# Cell type-specific inference of differential expression in spatial transcriptomics

**DOI:** 10.1101/2021.12.26.474183

**Authors:** Dylan M. Cable, Evan Murray, Vignesh Shanmugam, Simon Zhang, Michael Diao, Haiqi Chen, Evan Z. Macosko, Rafael A. Irizarry, Fei Chen

**Author notes:** These authors contributed equally (Rafael A. Irizarry, Fei Chen).

## Abstract

Spatial transcriptomics enables spatially resolved gene expression measurements at near single-cell resolution. There is a pressing need for computational tools to enable the detection of genes that are differentially expressed (DE) within specific cell types across tissue context. We show that current approaches cannot learn cell type-specific DE due to changes in cell type composition across space and the fact that measurement units often detect transcripts from more than one cell type. Here, we introduce a statistical method, Cell type-Specific Inference of Differential Expression (C-SIDE), that identifies cell type-specific patterns of differential gene expression while accounting for localization of other cell types. We model spatial transcriptomics gene expression as an additive mixture across cell types of general log-linear cell type-specific expression functions. This approach provides a unified framework for defining and identifying gene expression changes in a wide-range of relevant contexts: changes due to pathology, anatomical regions, physical proximity to specific cell types, and cellular microenvironment. Furthermore, our approach enables statistical inference across multiple samples and replicates when such data is available. We demonstrate, through simulations and validation experiments on Slide-seq and MER-FISH datasets, that our approach accurately identifies cell type-specific differential gene expression and provides valid uncertainty quantification. Lastly, we apply our method to characterize spatially-localized tissue changes in the context of disease. In an Alzheimer’s mouse model Slide-seq dataset, we identify plaque-dependent patterns of cellular immune activity. We also find a putative interaction between tumor cells and myeloid immune cells in a Slide-seq tumor dataset. We make our C-SIDE method publicly available as part of the open source R package https://github.com/dmcable/spacexr.

## Introduction

Spatial transcriptomics technologies profile gene expression in parallel across hundreds or thousands of genes across spatial measurement units, or *pixels* [1–9]. These technologies have the potential to associate gene expression with cellular context such as spatial position, proximity to pathology, or cell-to-cell interactions. Studying gene expression changes, termed *differential expression* (DE), within tissue context has the potential to provide insight into principles of organization of complex tissues and disorganization in disease and pathology [1, 10–13].

Current methods for addressing differential expression in spatial transcriptomics fall into two categories: nonparametric and parametric methods. Nonparametric differential expression methods [14–17] do not use constrained hypotheses about gene expression patterns, but rather fit general smooth spatial patterns of gene expression. Some of these approaches do not take cell types into account [14, 15], while others operate on individual cell types [17]. Discovering non-parametric differential gene expression can be advantageous in order to generate diverse exploratory hypotheses. However, if covariates are available, for example predefined anatomical regions, parametric approaches increase statistical power substantially and provide directly interpretable parameter estimates. Specific differential expression problems have been addressed with ad-hoc solutions such as detecting gene expression dependent on cell-to-cell colocalization [18] or anatomical regions [10, 19], but no general parametric framework is currently available. In contrast, general parametric frameworks have been widely applied across bulk and single-cell RNA-sequencing (scRNA-seq) to test for differences in gene expression across cell type, disease state, and developmental state, among other problems [20–22]. Furthermore, although multi-sample, multi-replicate differential expression methods exist for bulk and single-cell RNA-seq [20–22], no statistical framework accounting for technical and biological variation [23] across samples and replicates has been established for the spatial setting.

An important challenge unaddressed by current spatial transcriptomics DE methods is accounting for observations generated from cell type mixtures. In particular, sequencing-based, RNA-capture spatial transcriptomics technologies, such as Visium [7], DBiT-seq [6], GeoMx [8], and Slide-seq [1, 2], can capture multiple cell types on individual measurement pixels. The presence of cell type mixtures complicates the estimation of *cell type-specific differential expression* (i.e. DE within a cell type of interest) because different cell types have different gene expression profiles, independent of spatial location [24, 25]. Although imaging-based spatial transcriptomics technologies, such as MERFISH [3], seqFISH [5], ExSeq [9], and STARmap [4], have the potential to achieve single cell resolution, these technologies may encounter mixing or contamination across cell types due to diffusion or imperfect cellular segmentation [26]. Several methods [24, 27, 28] have been developed to identify cell type proportions in spatial transcriptomics datasets. However, at present no method accounts for cell type proportions in differential expression analysis. Here, we demonstrate how not accounting for cell type proportions leads to biased estimates of differential gene expression due to confounding caused by cell type proportion changes or contamination from other cell types.

In this work we introduce Cell type-Specific Inference of Differential Expression (C-SIDE), a general parametric statistical method that estimates cell type-specific differential expression in the context of cell type mixtures. The first step is to estimate cell type proportions on each pixel using a cell type-annotated single-cell RNA-seq (scRNA-seq) reference [24]. Next, we fit a parametric model, using predefined covariates such as spatial location or cellular microenvironment, that accounts for cell type differences to obtain cell type-specific differential expression estimates and corresponding standard errors. The model accounts for sampling noise, gene-specific overdispersion, multiple hypothesis testing, and platform effects between the scRNA-seq reference and the spatial data. Furthermore, when multiple experimental samples are available, the C-SIDE model permits statistical inference across multiple samples and/or replicates to achieve more stable estimates of population-level differential gene expression.

Using simulated and real spatial transcriptomics data, we show C-SIDE accurately estimates cell type-specific differential expression while controlling for changes in cell type proportions and contamination from other cell types. We also demonstrate how cell type mixture modelling increases power, especially when single cell type measurements are rare. Furthermore, on Slide-seq and MERFISH datasets, we demonstrate how C-SIDE’s general parametric framework enables testing differential gene expression for diverse hypotheses including spatial position or anatomical regions [29], cell-to-cell interactions, cellular environment, or proximity to pathology. By associating gene expression changes with particular cell types, we use C-SIDE to systematically link gene expression changes to cellular context in pathological tissues such as Alzheimer’s disease and cancer.

## Results

### Cell type-Specific Inference of Differential Expression learns cell type-specific differential gene expression in the context of spatial transcriptomics cell type mixtures

Here, we develop Cell type-Specific Inference of Differential Expression (C-SIDE), a statistical method for determining differential expression (DE) in spatial transcriptomics datasets (Figure 1a). C-SIDE inputs one or more experimental samples of spatial transcriptomics data, consisting of *Y_i,j,g_* as the observed RNA counts for pixel *i*, gene *j*, and experimental sample *g*. We then assume Poisson sampling so that,

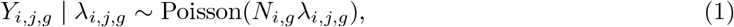

with *λ_i,j,g_* the expected count and *N_i,g_* the total transcript count (e.g. total UMIs) for pixel *i* on experimental sample *g*. Accounting for platform effects and other sources of technical and natural variability, we assume *λ_i,j,g_* is a mixture of *K* cell type expression profiles, defined by,

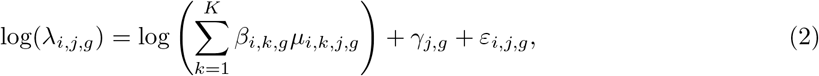

with *μ_i,k,j,g_* the cell type-specific expected gene expression rate for pixel *i*, gene *j*, experimental sample *g*, and cell type *k*; *β_i,k,g_* the proportion of cell type *k* contained in pixel *i* for experimental sample *g*; *γ_j,g_* a gene-specific random effect that accounts for platform variability; and *ε_i,j,g_* a random effect to account for gene-specific overdispersion.

**Figure 1:**
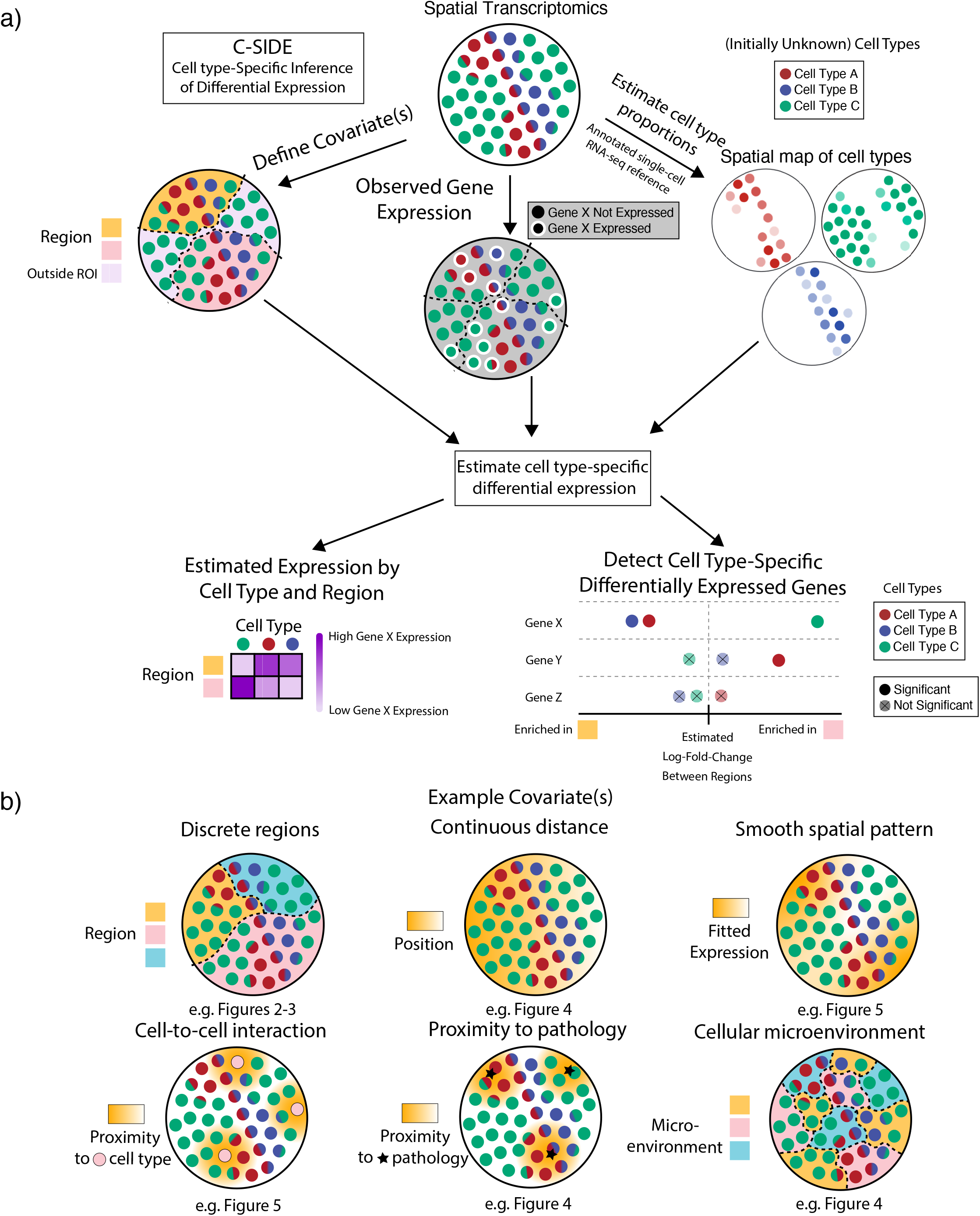
Cell type-Specific Inference of Differential Expression learns cell type-specific differential expression from spatial transcriptomics data. (a) Schematic of the C-SIDE Method. Top: C-SIDE inputs: a spatial transcriptomics dataset with observed gene expression (potentially containing cell type mixtures) and a covariate for differential expression. Middle: C-SIDE first assigns cell types to the spatial transcriptomics dataset, and covariates are defined. Bottom: C-SIDE estimates cell type-specific gene expression along the covariate axes. (b) Example covariates for explaining differential expression with C-SIDE. Top: Segmentation into multiple regions, continuous distance from some feature, or general smooth patterns (nonparametric). Bottom: density of interaction with another cell type or pathological feature or a discrete covariate representing the cellular microenvironment.

To account for cell type-specific differential expression, we model across pixel locations the log of the cell type-specific profiles *μ_i,k,j,g_* as a linear combination of *L* covariates used to explain differential expression. Specifically, we assume that,

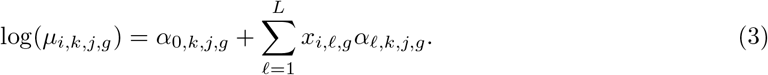

Here, *α*_0,*k*,*j*,*g*_ represents the intercept term for gene *j* and cell type *k* in sample *g*, and *x*_1,*ℓ*,*g*_ represents the *ℓ*’th *covariate*, evaluated at pixel *i* in sample *g*. Similarly as in linear and generalized linear models [30], *x*, also called the *design matrix*, represents predefined covariate(s) that explain differential expression, and the corresponding coefficients) *α_ℓ,k,j,g_* each represent the DE effect size of covariate *ℓ* for gene *j* in cell type *k* for sample *g*.

With this general framework we can describe any type of differential expression that can be parameterized with a log-linear model. Examples include (Figure 1b):

1. Differential expression between multiple regions. In this case, the tissue is manually segmented into multiple regions (e.g. nodular and anterior cerebellum, Figure 3). Design matrix *x* contains discrete categorical indicator variables representing membership in 2 or greater regions.
2. Differential expression due to cellular environment or state (special case of (1)). Pixels are discretely classified into local environments based on the surrounding cells (e.g. stages in the testes Slide-seq dataset, Figure 4).
3. Differential expression as a function of distance to a specific anatomical feature. In this case, *x* is defined as the spatial position or distance to some feature (e.g. distance to midline in the hypothalamus MERFISH dataset, Figure 4).
4. Cell-to-cell interactions. In this case, we define a cell-to-cell interaction as differential expression within one cell type (*A*) due to co-localization with a second cell type (*B*) (e.g. immune cell density in cancer, Figure 5). For this problem, *x* is the continuous density of cell type *B*.
5. Proximity to pathology. Similar to (4), except covariate *x* represents density of a pathological feature (e.g. Alzheimer’s A*β* plaque, Figure 4), rather than cell type density.
6. General spatial patterns (termed *nonparametric)*. In this case, we define design matrix *x* to be smooth basis functions [31], where linear combinations of these basis functions represent the overall smooth gene expression function and can accommodate any smooth spatial pattern.

**Figure 2:**
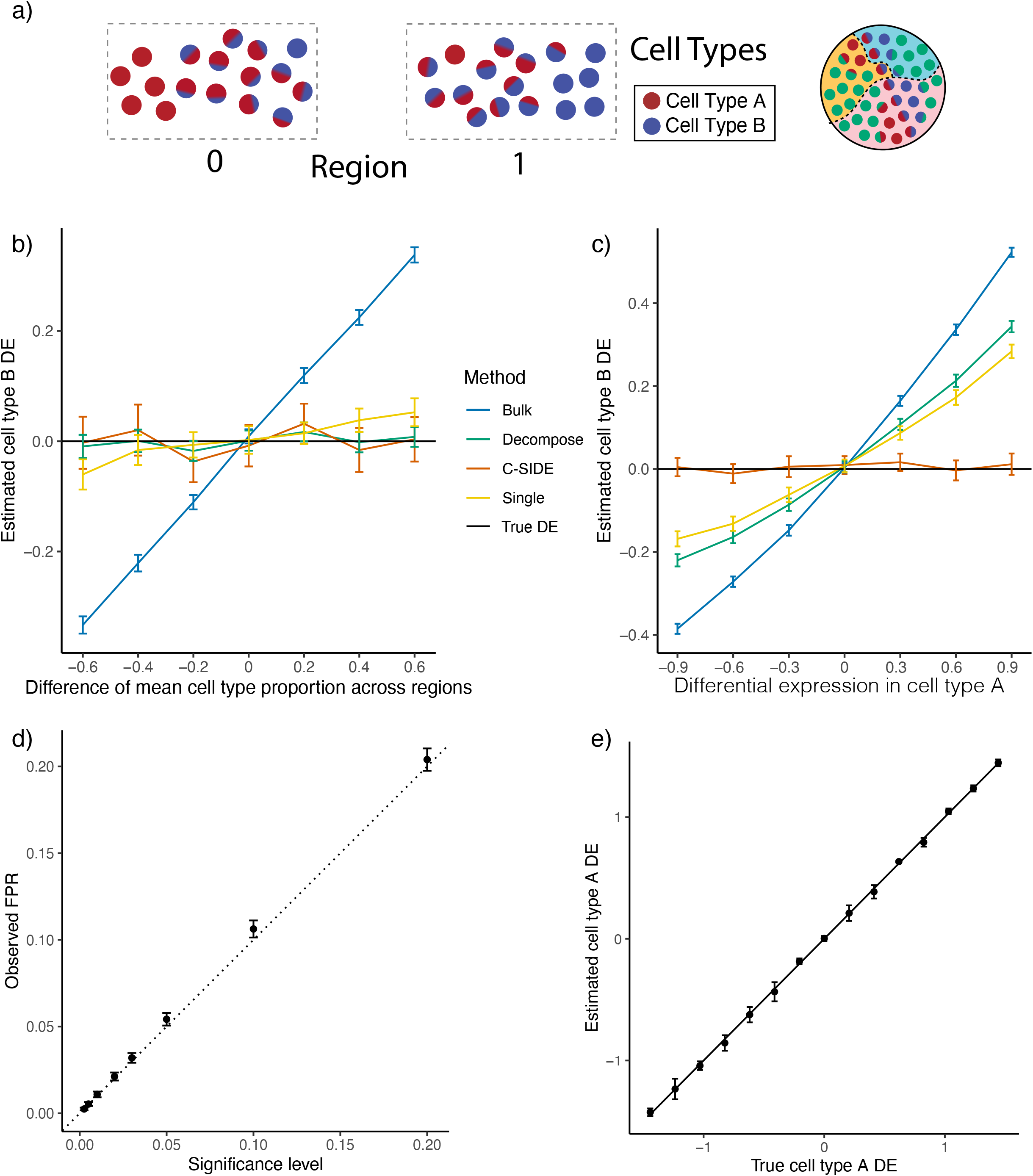
C-SIDE provides unbiased estimates of cell type-specific differential expression in simulated data. All: C-SIDE was tested on a dataset of simulated mixtures of single cells from a single-nucleus RNA-seq cerebellum dataset. Differential expression (DE) axes represent DE in log2-space of region 1 w.r.t. region 0. (a) Pixels are grouped into two regions, and genes are simulated with ground truth DE across regions. Each region contains pixels containing mixtures of various proportions between cell type A and cell type B. The difference in average cell type proportion across regions is varied across simulation conditions. (b) Mean estimated cell type B *Astn2* DE (differential expression) across two regions as a function of the difference in mean cell type proportion across regions. *Astn2* is simulated with ground truth 0 spatial DE, and an average of (*n* = 100) estimates is shown, along with standard errors. Black line represents ground truth 0 DE (cell type B). Four methods are shown: *Bulk*, *Decompose*, *Single*, and *C-SIDE* (see *Methods* for details). (c) Same as (b) for *Nrxn3* cell type B differential gene expression as a function of DE in cell type A, where *Nrxn3* is simulated to have DE within cell type A but no DE in cell type B. (d) For each significance level, C-SIDE’s false positive rate (FPR), along with ground truth identity line (s.e. shown, *n* = 1500, 15 genes, 100 replicates per gene). (e) C-SIDE mean estimated cell type A differential expression vs. true cell type A differential expression (average over *n* = 500 replicates, s.e. shown). Ground truth identity line is shown, and one gene is used for the simulation per DE condition (out of 15 total genes).

**Figure 3:**
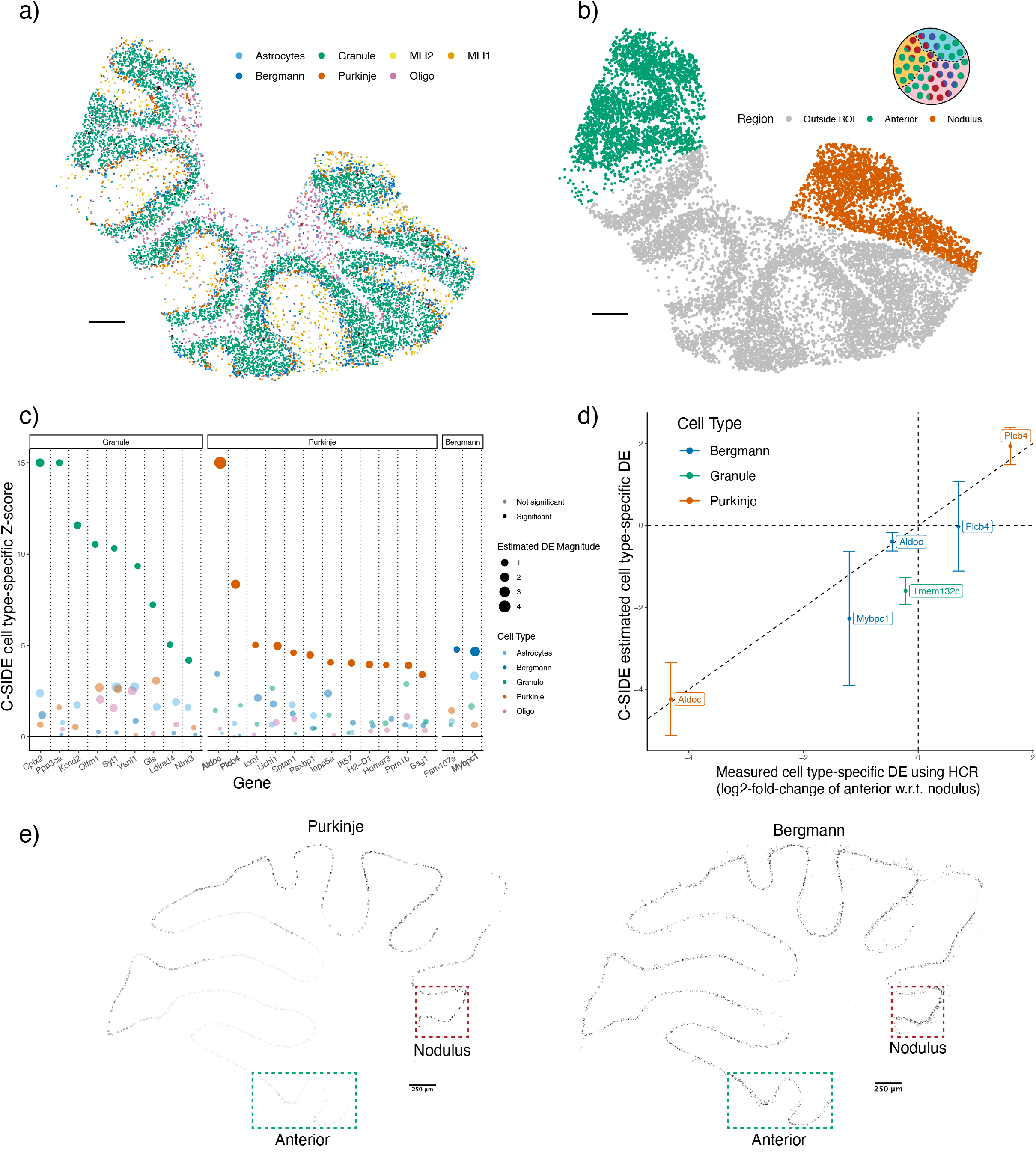
C-SIDE’s estimated cell type-specific differential expression is validated by HCR-FISH. (a) C-SIDE’s spatial map of cell type assignments in the cerebellum Slide-seq dataset. Out of 19 cell types, the seven most common appear in the legend. Reproduced from [24]. Three total replicates were used to fit C-SIDE. (b) Covariate used for C-SIDE, representing the anterior lobule region (green) and nodulus (red). Schematic refers to the C-SIDE problem type outlined in Figure 1b. (c) C-SIDE Z-score for testing for DE for each gene and for each cell type. Genes are grouped by cell type with maximum estimated DE, and estimated DE magnitude appears as size of the points. Bold genes appear below in HCR validation. (d) Scatterplot of C-SIDE DE estimates vs. HCR measurements for cell type-specific log2 differential expression. Positive values indicate gene expression enrichment in the anterior region. Error bars represent C-SIDE confidence intervals for predicted DE on a new biological replicate. A dotted identity line is shown, and cell types are colored. (e) HCR images of *Aldoc* continuous gene expression. Only pixels with high cell type marker measure-ments for Purkinje (left) and Bergmann (right) are shown. Regions of interest (ROIs) of nodulus and anterior regions are outlined in green and red, respectively. All scale bars 250 microns.

**Figure 4:**
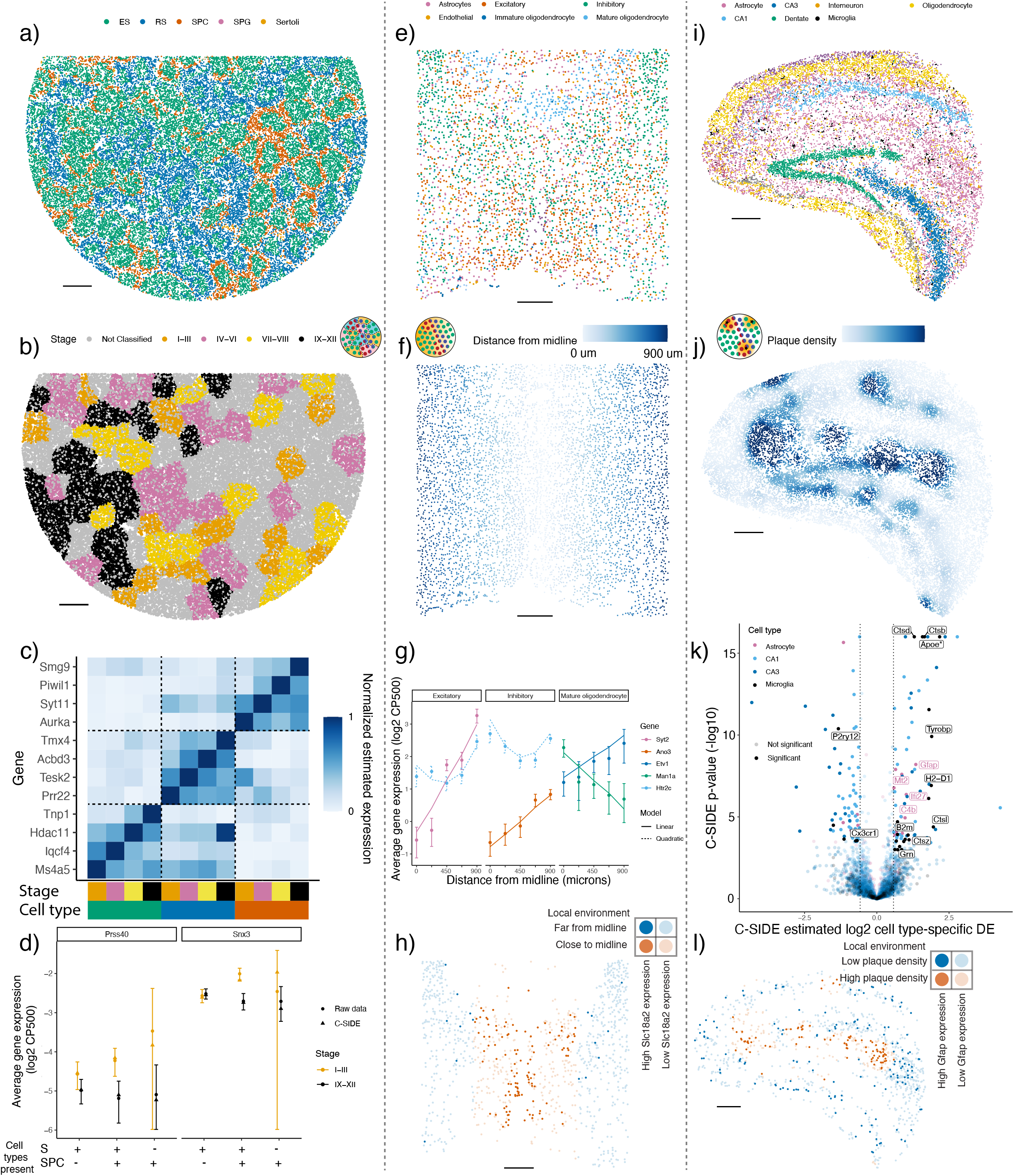
C-SIDE discovers cell type-specific differential expression in a diverse set of problems on testes, Alzheimer’s hippocampus, and hypothalamus datasets. All panels: results of C-SIDE on the Slide-seqV2 testes (left column), MERFISH hypothalamus (middle column), and Slide-seqV2 Alzheimer’s hippocampus (right column). Schematics in b,f,j reference C-SIDE problem types (Figure 1b). (a) C-SIDE’s spatial map of cell type assignments in testes. All cell types are shown, and the most common cell types appear in the legend. (b) Covariate used for C-SIDE in testes: four discrete tubule stages. (c) Cell type and tubule stage-specific genes identified by C-SIDE. C-SIDE estimated expression is standardized between 0 and 1 for each gene. Columns represent C-SIDE estimates for each cell type and tubule stage. (d) Log2 average expression (in counts per 500 (CP500)) of pixels grouped based on tubule stage and presence or absence of spermatid (S) cell types (defined as elongating spermatid (ES) or round spermatid (RS)) and/or spermatocyte (SPC) cell type. Circles represent raw data averages while triangles represent C-SIDE predictions, and error bars around circular points represent ± 1.96 s.d. (*Supplementary methods*). Genes *Prss40* and *Snx3* are shown on left and right, respectively. (e) Same as (a) for hypothalamus. (f) Covariate used for C-SIDE in hypothalamus: continuous distance from midline. (g) Log2 average expression (in counts per 500 (CP500)) of genes identified to be significantly differentially expressed by C-SIDE for each of the excitatory, inhibitory, and mature oligodendrocyte cell types. Single cell type pixels are binned according to distance from midline, and points represent raw data averages while lines represents C-SIDE predictions and error bars around points represent ± 1.96 s.d. (*Supplementary methods*). (h) Spatial visualization of *Slc18a2*, whose expression within inhibitory neurons was identified by C-SIDE to depend on midline distance. Red/blue represents inhibitory neurons close/far to midline, respectively. Bold points inhibitory neurons expressing *Slc18a2* at a level of at least 10 counts per 500. (i) Same as (a) for Alzheimer’s hippocampus, where four total replicates were used to fit C-SIDE. (j) Covariate used for C-SIDE in Alzheimer’s hippocampus: continuous density of beta-amyloid (A*β*) plaque. (k) Volcano plot of C-SIDE differential expression results in log2-space, with positive values corresponding to plaque-upregulated genes. Color represents cell type, and a subset of significant genes are labeled. Dotted lines represents 1.5x fold-change cutoff used for C-SIDE. (*): *Apoe* didn’t pass default C-SIDE gene filters(*Methods*) because 4x higher expression in astrocytes than microglia. (l) Spatial visualization of *Gfap*, whose expression within astrocytes was identified by C-SIDE to depend on plaque density. Red/blue represents the astrocytes in high/low plaque density areas, respectively. Bold points represent astrocytes expressing *Gfap* at a level of at least 1 count per 500. All scale bars 250 microns.

**Figure 5:**
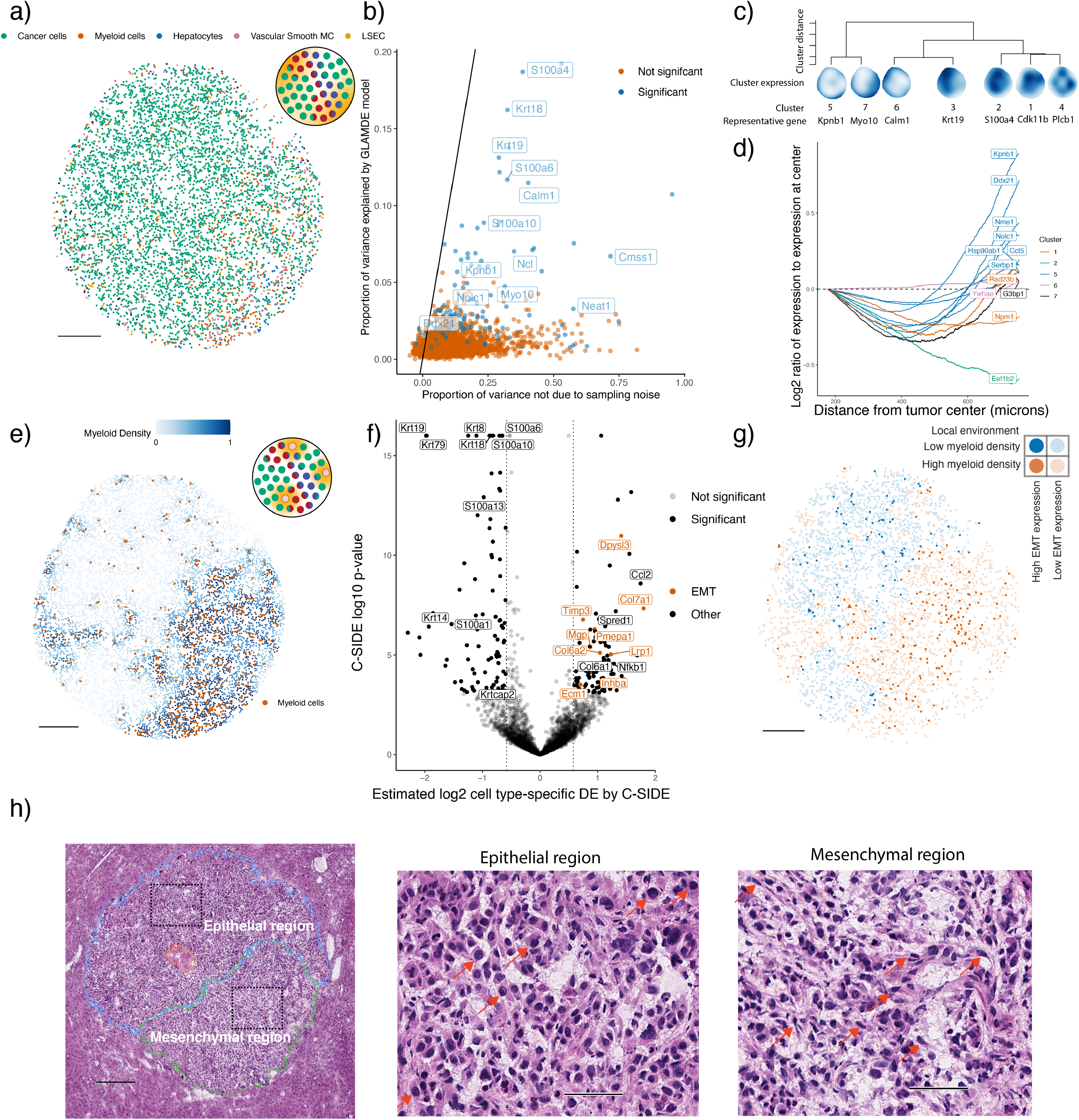
C-SIDE enables the discovery of differentially expressed pathways in a *Kras*^G12D/+^ *Trp53*^-/-^ (KP) mouse model. All panels: C-SIDE was run on multiple cell types, but plots represent C-SIDE results on the tumor cell type.Nonparametric C-SIDE results are shown in panels b–d, while parametric C-SIDE results are shown in panels e–h. (a) C-SIDE’s spatial map of cell type assignments. Out of 14 cell types, the five most common appear in the legend. (b) Scatter plot of C-SIDE *R*^2^ and overdispersion (defined as proportion of variance not due to sampling noise) for nonparametric C-SIDE results on the tumor cell type. Identity line is shown, representing the maximum possible variance that could be explained by any model. (c) Dendrogram of hierarchical clustering of (n = 162 significant genes) C-SIDE’s fitted smooth spatial patterns at the resolution of 7 clusters. Each spatial plot represents the average fitted gene expression patterns over the genes in each cluster. (d) Moving average plot of C-SIDE fitted gene expression (normalized to expression at center) as a function of distance from the center of the tumor for 12 genes in the *Myc* targets pathway identified to be significantly spatially differentially expressed by C-SIDE. (e) Covariate used for parametric C-SIDE: continuous density of myeloid cell types in the tumor. Schematic refers to C-SIDE problem type (Figure 1b). (f) Volcano plot of C-SIDE log2 differential expression results on the tumor cell type with positive values representing upregulation in the presence of myeloid immune cells. A subset of significant genes are labeled, and dotted lines represent 1.5x fold-change cutoff. (g) Spatial plot of total expression in tumor cells of the 9 differentially expressed epithelial-mesenchymal transition (EMT) genes identified by C-SIDE in (f). Red represents the tumor cells in high myeloid density areas, whereas blue represents tumor cells in regions of low myeloid density. Bold points represent tumor cells expressing these EMT genes at a level of at least 2.5 counts per 500. (h) Hematoxylin and eosin (H&E) image of adjacent section of the tumor. Left: mesenchymal (green), necrosis (red), and epithelial (blue) annotated tumor regions, with dotted boxes representing epithelial and mesenchymal areas of focus for the other two panels. Middle/right: enlarged images of epithelial (middle) or mesenchymal (right) regions. Red arrows point to example tumor cells with epithelial (middle) or mesenchymal (right) morphology. All scale bars 250 microns, except for (h) middle/right, which has 50 micron scale bars.

To estimate this complex model with a computationally tractable algorithm, we note that the gene expression variability across cell types is large enough that, in the first step, we can assume *μ_i,k,j,g_* does not vary with *i* and *g* and estimate *β* using a previously published algorithm [24]. Here, some pixels are identified as single cell types while others as mixtures of multiple cell types. Fixing the *β* estimates, we next use maximum likelihood estimation to estimate the cell type-specific DE coefficients *α* with corresponding standard errors, allowing for false discovery rate-controlled hypotheses testing (see *Methods* for details). Lastly, C-SIDE performs statistical inference across multiple replicates and/or samples, accounting for biological and technical variation across replicates, to estimate consensus population-level differential expression (*Methods*, Supplementary Figure 1).

Because ground truth cell type-specific DE is unknown in spatial transcriptomics data, we first benchmarked C-SIDE’s performance on a simulated spatial transcriptomics dataset in which gene expression varied across two regions. Considering the challenging situation where two cell types, termed *cell type A* and *cell type B*, are colocalized on pixels within a tissue, we simulated, using a single-nucleus RNA-seq cerebellum dataset, spatial transcriptomics mixture pixels with known proportions of single cells from two cell types known to spatially colocalize [32] (*Methods*, Figure 2a). Across two spatially-defined regions, we varied both the true cell type-specific gene expression of cell types A and B as well as the average cell type proportions of cell types A and B (Figure 2a, Supplementary Figure 2). We compared C-SIDE against three alternative methods (see *Methods* for details): *Bulk*, bulk differential expression (ignoring cell type); *Single*, single cell differential expression that approximates each cell type mixture as a single cell type; and *Decompose*, a method that decomposes mixtures into single cell types prior to computing differential expression. By varying cell type frequencies between the two regions without introducing differential expression, we observed that C-SIDE correctly attributes gene expression differences across regions to differences in cell type proportions rather than spatial differential expression (Figure 2b, Supplementary Figure 2); in contrast, the Bulk method incorrectly predicts spatial differential expression since it does not control for differences of cell type proportions across regions.

Next, we simulated cell type-specific differential expression (DE) by varying the differential expression in cell type A while keeping cell type B constant across regions. Background DE in cell type A contaminated estimates of differential expression in cell type B for all three alternatives models Bulk, Decompose, and Single (Figure 2c, Supplementary Figure 2). In contrast, C-SIDE’s joint model of cell type mixtures and cell type-specific differential expression correctly identified differential expression in cell type A, but not cell type B. Next, we verified that, under the null hypothesis of zero differential expression, C-SIDE’s false positive rate was accurately controlled, standard errors were accurately estimated, and confidence intervals contained the ground truth DE (Figure 2d, Supplementary Figure 2). Finally, when nonzero differential expression was simulated, C-SIDE achieved unbiased estimation of cell type-specific differential expression (Figure 2e). We also found that the power of C-SIDE depends on gene expression level, number of cells, and differential expression magnitude (Supplementary Figure 2). Thus, our simulations validate C-SIDE’s ability to accurately estimate and test for cell type-specific differential expression in the cases of asymmetric cell type proportions and contamination from other cell types.

### C-SIDE accurately identifies cell type-specific differential expression in spatial transcriptomics data

To validate C-SIDE’s ability to discover cell type-specific differential expression on spatial transcriptomics data, we collected Slide-seqV2 data [2] (including one replicate sourced from a prior study [24]) for three cerebellum replicates. We identified a spatial map of cell types (Figure 3a), previously shown to correspond to known cerebellum spatial architecture [24]. We used discrete localization in the anterior lobule or nodulus regions (Figure 3b), a known axis of spatial gene expression variation within the cerebellum [32], as a covariate and estimated cell type-specific DE across regions using C-SIDE (Figure 3c, Supplementary Figure 3, Supplementary Table 1). As experimental validation, we performed hybridization chain reaction (HCR) on four genes identified to be differentially expressed in specific cell types, and we observed high correspondence between C-SIDE’s estimates of cell type-specific differential expression and DE measurements from HCR data (Figure 3d, *R*^2^ = 0.89). For example, we examined *Aldoc* and *Plcb4*, two genes expressed in both Purkinje and Bergmann cell types, which are known to spatially colocalize in the cerebellum and appear as mixtures on Slide-seq pixels [24]. C-SIDE determined that both *Aldoc* (log2-fold-change = −4.24, p < 10^-8^) and *Plcb4* (log2-fold-change = 1.93, p < 10^-8^) were differentially expressed in the Purkinje cell type, but not the Bergmann cell type. Similarly, HCR images of *Aldoc* and *Plcb4* showed substantial differential expression within Purkinje cells across the nodulus and anterior lobule, whereas expression within Bergmann cells was relatively even across regions (Figure 3d–e). We conclude that C-SIDE can successfully identify cell type-specific spatial differential expression in spatial transcriptomics tissues, even when multiple cell types are spatially colocalized.

### C-SIDE solves a diverse array of differential gene expression problems in spatial transcriptomics

We next explored the effect of discrete cellular microenvironments on cell type-specific DE in the mouse testes Slide-seq dataset [12]. C-SIDE’s testes principal cell type assignments (Figure 4a) revealed tubular structures corresponding to cross-sectional sampling of seminiferous tubules. Individual tubules have distinct stages of spermatogonia development, grouped into four classes of stages I–III, IV–VI, VII–VIII, and IX–XII, which were determined from the prior testes Slide-seq study using tubule-level gene expression clustering [12] (Figure 4b). Given that each tubule stage represents a distinct microenvironment along the testes developmental trajectory, we applied C-SIDE to identify genes that were differentially expressed, for each cell type, across tubule stages (Supplementary Table 2). Furthermore, C-SIDE identified genes expressed in a single tubule stage within a single cell type (Figure 4c) which are known drivers of cellular development across stages [12]. For instance, the gene *Tnpl* was identified by C-SIDE as upregulated in the IX–XII stage within the elongating spermatid (ES) cell type, in agreement with the known biological role of *Tnp1* in nuclear remodeling of elongating spermatids at the late tubule stage [33] (Supplementary Figure 4). After identifying stage-specific genes within each cell type, we additionally found that a majority of C-SIDE-identified stage-specific genes followed cyclic patterns across stages, consistent with previously-characterized cyclic gene regulation in what is referred to as the seminiferous epithelial cycle [34] (Supplementary Figure 4).

Next, we evaluated C-SIDE’s ability to identify DE for cell types that primarily appear as mixtures with other cell types, particularly the spermatocyte (SPC) cell type. According to C-SIDE cell type assignments, SPC frequently co-mixes with the ES and round spermatid (RS) cell types, consistent with the known colocalization of spermatocytes with spermatids from previous histological studies [35] (Supplementary Figure 4). Due to C-SIDE’s ability to learn DE from cell type mixtures, C-SIDE obtained increased power for identifying differentially expressed genes compared to a DE method that only uses single cell type pixels (see *Supplementary Methods* for details, Supplementary Figure 4), especially for spermatocyte cell type (217 significant SPC DE genes discovered by C-SIDE vs. 1 DE gene for the single cell method). In order to validate C-SIDE’s determination that *Prss40* (log2-fold-change = 1.72, p = 8 · 10^-5^) and *Snx3* (log2-fold-change = 1.17, p < 10^-8^) were differentially expressed, between stage I–III and stage IX–XII, specifically in the SPC cell type, we compared the average gene expression for three categories of testes pixels: pixels containing spermatid cell types, but not SPC (called S+, SPC-); pixels containing both spermatid and SPC cell types (S+, SPC+); and pixels containing SPC but not spermatids (S-, SPC+) (Figure 4d). For both genes, differential expression across stages was not observed in (S+, SPC-) pixels, indicating that the spermatid cell types do not exhibit DE. However, (S+, SPC+) pixels are significantly differentially expressed across stages, enabling C-SIDE to infer DE specifically in the SPC cell type. On the other hand, (S-, SPC+) pixels, which include SPC single cells, are not significantly differentially expressed across regions, due to their low sample size. Therefore, C-SIDE’s ability to handle cell type mixtures uniquely enables the discovery of differential expression, even in cell types that only appear as mixtures with other cell types.

### C-SIDE identifies spatial gene expression changes in imaging-based technologies

Next, we demonstrated the utility of C-SIDE on an imaging-based spatial transcriptomics dataset (i.e. MERFISH) which achieves closer to single-cell resolution compared to capture-based spatial transcriptomics technologies (e.g. Slide-seq, Visium), which contain frequent cell type mixtures [24]. To do so, we applied C-SIDE to a MERFISH dataset collected in the mouse hypothalamus. During development, hypothalamic progenitors create radial projections out from the hypothalamic midline, which are used as scaffolds for the migration of differentiating daughter cells [36]. Thus, we investigated radial distance to the hypothalamus midline as a predictor of differential expression in hypothalamus cell types. First, we assigned cell types and found them to be consistent with the prior MERFISH hypothalamus study [11] (Figure 4e). Although C-SIDE mostly assigned single cell types to MERFISH pixels, a non-negligible proportion (12.6% double cell type pixels out of *n* = 3790 total single and double cell type pixels) of pixels were assigned as mixtures of more than one cell type. Next, we computed midline distance as a covariate for C-SIDE (Figure 4f), and we next detected genes in hypothalamus excitatory, inhibitory, and mature oligodendrocyte cell types whose expression depended either linearly or quadratically on distance from the midline (Figure 4g, Supplementary Table 3–4). For instance, *Slc18a2* (Figure 4h), identified by C-SIDE as differentially upregulated within inhibitory neurons near the midline (log2-fold-change = 6.14, p < 10^-8^), is required for dopaminergic function in certain inhibitory neuronal subtypes [37], which are known to localize near the hypothalamus midline [11].

### C-SIDE enables discovery of A*β* plaque-dependent cell type-specific differential expression in Alzheimer’s disease

We next explored the use of pathological staining, in particular A*β* plaques, as a continuous covariate for cell type-specific gene expression changes. To do so, we performed Slide-seqV2 on the hippocampal region of a genetic mouse model of amyloidosis in Alzheimer’s disease (AD) [38] (J20, n= 4 slices, *Methods*). C-SIDE identified spatial maps of cell types (Figure 4i) which were consistent with past characterizations of hippocampus cellular localization [24]. We collected paired A*β* plaque staining images (Anti-Human A*β* Mouse IgG antibody, *Methods*) to quantify the A*β* plaque density to use as a covariate for C-SIDE (Figure 4j, Supplementary Figure 5). We then used C-SIDE to identify genes whose expression depended in a cell type-specific manner on A*β* plaque density (Figure 4k, Supplementary Table 5). For instance, we found that *Gfap* was enriched in astrocytes colocalizing with A*β* plaque (Figure 4l, Supplementary Figure 5, log2-fold-change = 1.35, p < 10^-8^), a result corroborated by studies that have established the role of *Gfap* in attenuating the proliferation of A*β* plaques [39]. C-SIDE additionally discovered upregulation in astrocytes of the *C4b* complement gene (log2-fold-change = .85, p = 1 · 10^-4^), which is involved in plaque-associated synaptic pruning in Alzheimer’s disease [40–42]. Moreover, several cathepsin proteases including *Ctsb* (log2-fold-change = 1.65, p < 10^-8^), *Ctsd* (log2-fold-change = 1.30, p < 10^-8^) *Ctsl* (log2-fold-change = 1.96, p = 4 · 10^-6^), and *Ctsz* (log2-fold-change = 1.11, p = 3 · 10^-4^) were determined to be differentially upregulated in microglia around plaque, consistent with prior evidence that cathepsins are involved with amyloid degradation in Alzheimer’s disease [43] (Supplementary Figure 5). In microglia, we also identified known homeostatic microglia markers [44–46] including *P2ry12* (log2-fold-change = −1.33, p < 10^-8^) and *Cx3cr1* (log2-fold-change = −0.68, p = 3 · 10^-4^) as downregulated in the presence of plaque. *Apoe*, which is known to have A*β* plaque-dependent upregulation within microglia [47], was also detected as significant (log2-fold-change = 1.58, p < 10^-8^), although it did not pass default C-SIDE gene filters (*Methods*) due to its four-fold higher expression in astrocytes than microglia. Finally, the anti-inflammatory gene *Grn* was determined by C-SIDE to be upregulated in microglia near plaque (log2-fold-change = 0.79, p = 6 · 10^-4^), consistent with prior knowledge [48].

### C-SIDE discovers tumor-immune signaling in a mouse tumor model

Finally, we applied C-SIDE to identify genes with cell type-specific spatial differential expression in a Slide-seq dataset of a *Kras*^G12D/+^ *Trp53*^-/-^ (KP) mouse tumor model [49, 50], where we analyzed a single metastatic lung adenocarcinoma tumor deposit in the liver. We first used C-SIDE to generate a spatial map of cell types and found several cell types within the tumor, including both tumor cells and myeloid cells (Figure 5a). Next, we ran C-SIDE nonparametrically to discover arbitrary smooth gene expression patterns (see *Supplementary Methods* for details, Supplementary Table 6). For gene expression within the tumor cell type, this procedure identified three categories of genes: genes with variable expression purely due to sampling noise rather than biology, genes exhibiting biological variation partially explained by the spatial C-SIDE model, and genes exhibiting biological variation not explained by the spatial model (Figure 5b, Supplementary Figure 6). We then hierarchically clustered the C-SIDE fitted spatial patterns of significant differentially expressed genes within the tumor cell type into seven clusters with distinct spatial patterns (Figure 5c, Supplementary Figure 6). We tested each cluster for gene set enrichment (see *Supplementary Methods* for details), and we identified the *Myc* targets gene set as enriched in cluster 5 (7 out of 12 genes, *p* =2 · 10^-4^, two-sided binomial test, Supplementary Table 7, 1 significant gene set out of 50 tested), a cluster with a spatial pattern of overexpression at the tumor boundary (Figure 5d). High expression of *Myc* target genes is potentially indicative of an increased rate of proliferation [51] at the boundary, which has been previously proposed as a correlate of tumor severity [52]. For example, the *Myc* target found to have the most differential upregulation at the tumor boundary, *Kpnb1* (Supplementary Figure 6, p = 1 · 10^-5^), has been previously been identified as an oncogene that drives cell proliferation and suppresses apoptosis [53, 54].

Given the substantial variation in tumor cell spatial expression patterns, we next tested if such variability could be explained by cell-to-cell interactions with immune cells, which have been shown to influence tumor cell behavior in prior studies [55–57]. Using myeloid cell type density as the C-SIDE covariate (Figure 5e), C-SIDE identified genes with immune cell density-dependent cell type-specific differential expression (Figure 5f, Supplementary Table 8), including several genes that were also discovered by our nonparametric procedure (Supplementary Figure 6). One of the genes with the largest effects, *Ccl2* (log2-fold-change = 1.74, p < 10^-8^), is a chemotactic signaling molecule known to attract myeloid cells [58, 59]. Furthermore, we tested C-SIDE’s DE gene estimates for aggregate effects across gene sets and found that the epithelial-mesenchymal transition (EMT) pathway was significantly upregulated on average near immune cells (Figure 5f, Supplementary Figure 6, p = 0.0011, permutation test (see *Methods*), 1 significant gene set out of 50 tested, Supplementary Table 7). C-SIDE additionally identified *Nfkb1* as upregulated in tumor cells in immune-rich regions (log2-fold-change = 1.10, p = 1 · 10^-5^), a gene that has been previously implicated in positively regulating the EMT pathway of tumor cells [60, 61]. Moreover, the majority of tumor cells exhibiting a mesenchymal phenotype were located in immune-rich regions (Figure 5g). Furthermore, morphological analysis and annotation of an hematoxylin and eosin (H&E) stained adjacent section of the tumor demonstrated a clear increase in the number of spindle-shaped tumor cells relative to polygonal-shaped tumor cells in the immune rich-areas (Figure 5h). The collective morphological and gene expression changes suggest a role for the immune microenvironment in influencing the epithelial-mesenchymal transition in this tumor model [62]. Therefore, both exploratory nonparametric C-SIDE and more targeted immune cell-dependent DE reveal biologically-relevant signatures of differential gene expression.

## Discussion

Elucidating spatial sources of differential gene expression is a critical challenge for understanding biological mechanisms and disease with spatial transcriptomics. Here we introduced C-SIDE, a statistical method to detect cell type-specific DE in spatial transcriptomics datasets. C-SIDE takes as input one or more biologically-relevant covariates, such as spatial position or cell type colocalization, and identifies genes, for each cell type, that significantly change their expression as a function of these covariates. Tested on simulated spatial transcriptomics data, C-SIDE obtained unbiased estimation of cell type-specific differential gene expression with a calibrated false positive rate, while other methods were biased from changes in cell type proportion or contamination from other cell types. In the cerebellum, we additionally used HCR experiments to validate C-SIDE’s ability to identify cell type-specific DE across regions. We further applied C-SIDE to a detect differential expression depending on tubular microenvironment in the testes, midline distance in the MERFISH hypothalamus, and A*β* plaque density in the Alzheimer’s model hippocampus. Finally, we applied both nonparametric and parametric C-SIDE procedures in a mouse tumor model to discover an increase in tumor cells undergoing EMT transition in immune-rich regions.

Several studies have established the importance of accounting for cell type mixtures in assigning cell types in spatial transcriptomics data [24, 27, 28]. However, it remains a challenge to incorporate cell type proportions into models of cell type-specific spatial differential gene expression. C-SIDE enables such cell type-specific DE discovery by creating a statistical model of cell type-specific differential gene expression in the presence of cell type mixtures. In this study, we demonstrated how other potential solutions, such as bulk DE, approximation as single cell types, and decomposition into single cell types can be confounded by cell type proportion changes and contamination from other cell types. C-SIDE solves these issues by controlling for cell type proportions and jointly considering differential expression within each cell type. Even in imaging-based spatial transcriptomics methods such as MERFISH that mostly contain single cell type pixels, we detected some pixels with cell type mixtures, indicating potential diffusion or imperfect cell segmentation [26]. To control for cell type proportions in DE analysis, C-SIDE can estimate cell types directly or import cell type proportions from any cell type mixture identification method [24, 27, 28].

C-SIDE provides a unified framework for detecting biologically-relevant differential expression in spatial transcriptomics tissues along diverse array of axes including spatial distance, proximity to pathology, cellular microenvironment, and cell-to-cell interactions. In settings without prior biological hypotheses, C-SIDE may be run nonparametrically to discover general cell type-specific spatial gene expression patterns. When using problem-specific knowledge to generate biologically-relevant DE predictors, parametric C-SIDE efficiently detects DE genes along the parametric hypothesis axes. C-SIDE can also be used to test among multiple models of differential expression, such as the linear and quadratic models applied to the hypothalamus dataset. C-SIDE can also utilize multiple covariates in a joint model of gene expression, such as spatial position and cell type colocalization, although more complicated models require more data to fit accurately. Beyond individual samples, C-SIDE can also perform differential expression statistical inference at the population level across multiple replicates or biological samples, including modeling biological and technical variability in complex multi-sample, multi-replicate experiments. Multi-replicate experiments, though more costly, produce more robust DE estimates by reducing spurious discoveries of DE on single replicates.

One challenge for C-SIDE is obtaining sufficient DE detection statistical power, which we observed can be hindered by low gene expression counts, small pixel number, or rare cell types. An advantage of C-SIDE is that it increases its statistical power by including cell type mixture pixels in its model. Ongoing technical improvements in spatial transcriptomics technologies [2] such as increased gene expression counts, higher spatial resolution, and increased pixel number, have the potential to dramatically increase the discovery rate of C-SIDE. Another limitation of C-SIDE is the requirement of an annotated single-cell reference for reference-based identification of cell types in the cell type assignment step. Although single-cell atlases are increasingly available for biological tissues, they may contain missing cell types or substantial platform effects [24], and certain spatial transcriptomics tissues may lack a corresponding single-cell reference.

We envision C-SIDE to be particularly powerful in the context of bridging cell type-specific gene expression changes in pathology. Here, we demonstrate this in two contexts: one, wherein we leverage histological features (A*β* plaques) as a covariate, and two, wherein we nominate tumor-immune interactions as a covariate. In the first, prior Alzheimer’s disease (AD) studies have discovered candidate genes for disease-relevance through GWAS [63], bulk RNA and protein differences between AD and control samples [64], and single cell expression differences of disease associated cellular subtypes [41]. Here, with C-SIDE, we identify many genes previously identified by these methods including *Gfap*in astrocytes [39] and *Apoe* in microglia [47]; furthermore, we take known disease-level associations a step further towards mechanistic understanding by directly associating spatial plaque localization with cell type-specific differential expression. For example, prior studies have established an association between complement pathway activation in plaque-dense areas with synaptic pruning [40] and neuronal degeneration [41] leading to cognitive decline. Using C-SIDE we provide evidence for the upregulation of complement protein *C4b* specifically within plaque-localized astrocytes [65]. Thus, amyloid plaques may trigger a cytokine-dependent signaling cascade that stimulates the expression of complement genes in astrocytes, as supported by prior studies [42]. In contrast to *C4b* upregulation, homeostatic microglia marker *P2ry12*, discovered by C-SIDE to be negatively plaque-associated within microglia, has been shown to be downregulated in microglia in Alzheimer’s disease (AD), a phenomena associated with neuronal cell loss [44]. *P2ry12* is involved in early stage nucleotide-dependent activation of microglia and is downregulated in later stages of activated microglia [46]. We hypothesize that plaque-dense areas in AD trigger microglia activation which downregulates homeostatic microglia genes such as *P2ry12*. Lastly, the granulin gene (Grn), discovered by C-SIDE as upregulated in microglia near plaques, is an anti-inflammatory gene that attenuates microglia activation [66]. It has been shown to be upregulated in plaque-localized microglia in AD [48] and to potentially have a role in reducing plaque deposition and cognitive pathological effects in AD [67] and other pathological protein aggregates [68].

Second, C-SIDE has the potential to elucidate tissue interactions driving system-level behavior in complex tissues. For example, recent studies have characterized cell-to-cell interactions of immune cells influencing the behavior of tumor cells [55–57]. Consistent with these studies, on a Slide-seq dataset of a mouse tumor model, C-SIDE identified several genes whose expression within tumor cells was upregulated near myeloid immune cells. We postulate that the tumor cells and myeloid cells are involved in a synergistic feedback loop, driven by cell-to-cell signaling. For example, *Ccl2*, found by C-SIDE to be upregulated in immune-adjacent tumor cells, is known to chemotactically recruit myeloid cells and to induce pro-tumorigenic behavior, including growth, angiogenesis, and metastasis, in myeloid cells [58, 59]. Another synergistic immune-tumor interaction identified by C-SIDE is the myeloid-associated upregulation of the epithelial-mesenchymal transition (EMT) pathway, known to be involved in tumor development and metastasis [62]. Although C-SIDE established an association between immune cell colocalization and mesenchymal-like tumor cell state, conclusive establishment of mechanism of causation requires future experimentation. Among other hypotheses, it is plausible that myeloid cells induce tumor cells to undergo the EMT transition, potentially through the *NF-κB* (also identified as upregulated by C-SIDE) signaling pathway, as supported by other studies [55–57, 62]. Future work is necessary to characterize this phenomena across a broader cohort of samples and to establish specific molecular mechanisms. Overall, these results highlight the power of combining the C-SIDE framework with pathological measurements to understand cell type-specific responses to disease and injury. We envision C-SIDE as a powerful framework for the systematic study of the impacts of spatial and environmental context on cellular gene expression in spatial transcriptomics data.

## Methods

### C-SIDE model

Here, we describe Cell type-Specific Inference of Differential Expression (C-SIDE), a statistical method for identifying differential expression (DE) in spatial transcriptomics data. Please first refer to the overall definition of the C-SIDE model in equations (1), (2), and (3). Prior to fitting C-SIDE, the *design matrix x* is predefined to contain *covariates*, variables on which gene expression is hypothesized to depend such as spatial position or cellular microenvironment. Recall that *x_i,ℓ,g_* represents the *ℓ*’th covariate, evaluated at pixel *i* in experimental sample *g*. For each covariate *x*.*_ℓ,g_*, there is a corresponding coefficient *α_ℓ,k,j,g_*, representing a gene expression change across pixels per unit change of *x*.,_*ℓ,g*_ within cell type *k* of experimental sample *g*. Next, recall from (2) random effects *γ_j,g_* and *ε_i,j,g_*, which we assume both follow normal distributions with mean 0 and standard deviations *σ_γ,g_* and *σ_ε,j,g_*, respectively. We designed the overdispersion magnitude, *σ_εj,g_* to depend on gene *j* because we found evidence that the overdispersion depends on gene *j* (Supplementary Figure 7), and modeling gene-specific overdispersion is necessary for controlling the false-positive rate of C-SIDE.

Due to our finding that genes can exhibit DE in some but not all cell types (see e.g. Figure 3c), C-SIDE generally does not assume that genes share DE patterns across cell types, allowing for the discovery of cell type-specific DE. We also developed an option where DE can be assumed to be shared across cell types (*Supplementary Methods*). We note that C-SIDE can be thought of as a modification of the generalized linear model (GLM) [30] in which each cell type follows a cell type-specific log-linear model before an additive mixture of all cell types is observed. See Fitting the C-SIDE modeland Hypothesis testingfor C-SIDE model fitting and hypothesis testing, respectively.

### Parameterization of the design matrix

For specific construction of design matrix *x* for each dataset, see *Cell type estimation and construction of covariates*. Recall the specific examples of design matrix *x* presented in Figure 1b. In general, we note that *x* can take on the following numerical forms:

1. Indicator variable. In this case, *x_i,ℓ,g_* is always either 0 or 1. This represents differential expression due to membership within a certain spatially-defined pixel set of interest. The coefficient *α_k,j,g_* is interpreted as the log-ratio of gene expression between the two sets for cell type *k* and gene *j* in experimental sample *g*.
2. Continuous variable. In this case, *x_i,ℓ,g_* can take on continuous values representing, for example, distance from some feature or density of some element. The coefficient *α_ℓ,k,j,g_* is interpreted as the log-fold-change of gene expression per unit change in *x_i,ℓ,g_* for cell type *k* and gene *j* in sample *g*.
3. Multiple categories. In this case, we use *x* to encode membership to finitely many *L* ≥ 2 sets. For each 1 ≤ *ℓ* ≤ *L*, we define *x_i,ℓ,g_* to be an indicator variable representing membership in set *ℓ* for sample *g*. To achieve identifiability, the intercept is removed. The coefficient *α_ℓ,k,j,g_* is interpreted as the average gene expression in set *ℓ* for cell type *k* and gene *j*. Cell type-specific differential expression is determined by detecting changes in *α_ℓ,k,j,g_* across *ℓ* within cell type *k* and sample *g*.
4. Nonparametric. In this case, we use *x* to represent *L* smooth basis functions, where linear combinations of these basis functions represent the overall smooth gene expression function. By default, we use thin plate spline basis functions, calculated using the `mgcv` package [31].

In all cases, we normalize each *x_i,ℓ,g_* to range between 0 and 1. The problem is equivalent under linear transformations of *x*, but this normalization helps with computational performance. The intercept term, when used, is represented in *x* as a column of 1’s.

### Fitting the C-SIDE model

C-SIDE estimates the parameters of (1), (2), and (3) via maximum likelihood estimation. First, we note that all parameters and parameter relationships in the model are independent across samples, so we fit the model independently for each sample. We will return to the issue of population inference across multiple samples in Statistical inference on multiple samples/replicates. Next, the parameters of *β_i,k_* and *γ_j_* are estimated by the RCTD algorithm as previously described [24]. We note that C-SIDE can also optionally import cell type proportions from external cell type proportion identification methods [27, 28]. Here, some pixels are identified as single cell types while others as mixtures of multiple cell types. We can accurately estimate cell type proportions and platform effects without being aware of differential spatial gene expression because differential spatial gene expression is smaller than gene expression differences across cell types. After determining cell type proportions, C-SIDE estimates gene-specific overdispersion magnitude *σ_ε,j,g_* for each gene by maximum likelihood estimation (see *Supplementary Methods* for details). Finally, C-SIDE estimates the DE coefficients *α* by maximum likelihood estimation. For the final key step of estimating *α*, we use plugin estimates (denoted by ^) of *β*, *γ*, and *σ_ε_*. After we substitute (3) into (1) and (2), we obtain:

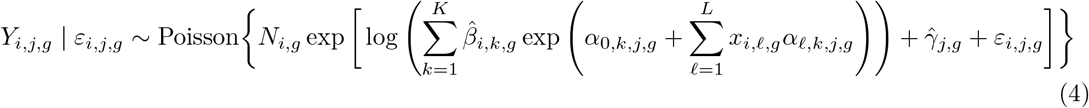

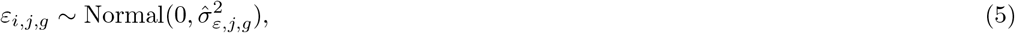

We provide an algorithm for computing the maximum likelihood estimator of *α*, presented in the *Supplementary Methods*. Our likelihood optimization algorithm is a second-order, trust-region based optimization (see *Supplementary Methods* for details). In brief, we iteratively solve quadratic approximations of the log-likelihood, adaptively constraining the maximum parameter change at each step. Critically, the likelihood is independent for each gene *j* (and sample *g*), so separate genes are run in parallel in which case there are *K* × (*L* +1) *α* parameters per gene and sample.

### Hypothesis testing

In addition to estimating the vector *α_j,g_* (dimensions *L* + 1 by *K*) for gene *j* and sample *g*, we can compute standard errors around *α_j,g_*. By asymptotic normality (see *Supplementary Methods* for details), we have approximately that (setting *n* to be the total number of pixels),

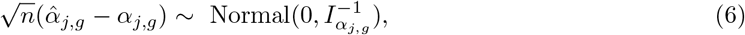

where *I_α,j,g_* is the Fisher information of model (4), which is computed in the *Supplementary Methods*. Given this result, we can compute standard errors, confidence intervals, and hypothesis tests. As a consequence of (6), the standard error of *α_ℓ,k,j,g_*, denoted *s_ℓ,k,j,g_*, is 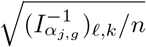.

First, we consider the case where we are interested in a single parameter, *α_ℓ,k,j,g_*, for *ℓ* and *g* fixed and for each cell type *k* and gene *j*; for example, *α_ℓk,j,g_* could represent the log-fold-change between two discrete regions. In this case, for each gene *j*, we compute the z-statistic, 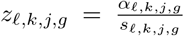. Using a two-tailed z-test, we compute a *p*–value for the null hypothesis that *α_ℓ,k,j,g_* = 0 as *p_ℓ,k,j,g_* = 2 * *F*(–|*z_ℓ,k,j,g_*|), where *F* is the distribution function of the standard Normal distribution. Finally, q-values are calculated across all genes within a cell type in order to control the false discovery rate using the Benjamini-Hochberg procedure [69]. We used a false discovery rate (FDR) of .01 (0.1 for nonparametric case) and a fold-change cutoff of 1.5 (N/A for nonparametric case). Additionally, for each cell type, genes were pre-filtered so that the expression within the cell type of interest had a total expression of at least 15 unique molecular identifiers (UMIs) over all pixels and at least 50% as large mean normalized expression as the expression within each other cell type.

For the multi-region case, we instead test for differences of pairs of parameters representing the average expression within each region. As a result, *p*–values are scaled up due to multiple hypothesis testing. We select genes which have significant differences between at least one pair of regions. For other cases in which we are interested in multiple parameters, for example the nonparametric case, we test each parameter individually and scale *p*-values due to multiple hypothesis testing.

### Statistical inference on multiple samples/replicates

C-SIDE can be run on either one or multiple biological replicates and/or samples. In the case of multiple replicates, we recall *α_g_* and *s_g_* are the differential expression and standard error for replicate *g*, where 1 ≤ *g* ≤ *G*, and *G* > 1 is the total number of replicates. We now consider testing for differential expression across all replicates for covariate *ℓ*, cell type *k*, and gene *j*. In this case, we assume that additional biological or technical variation across samples exists, such that each unknown *α_g_* is normally distributed around a population-level differential expression *A*, with standard deviation *τ*:

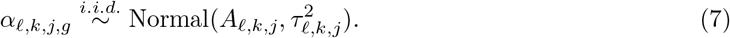

Under this assumption, and using (6) for the distribution of the observed single-sample estimates 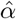, we derive the following feasible generalized least squares estimator of *A* (see *Supplementary Methods* for details),

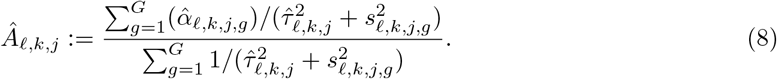

Here, 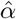 and *s* are obtained from C-SIDE estimates on individual samples (see (6)), whereas 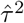 represents the estimated variance across samples (Supplementary Figure 7). Please see the *Supplementary Methods* for additional details such as the method of moments procedure [70] for estimating 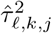 and the standard errors of *A*. Intuitively, our estimate of the population-level differential expression is a variance-weighted sum over the DE estimates of individual replicates, and we note that our multiple-replicate approach is similar to widely used meta-analysis methods [70, 71]. As we have obtained estimates and standard errors of *A*, these are subsequently used in hypothesis testing for the hypothesis that *A_ℓ,k,j_* = 0 in a manner identical to what is described above in Hypothesis testing for the single replicate case. We also derived a version of this estimator for the case where there are multiple biological samples and multiple replicates within each sample (*Supplementary Methods*).

### Collection and preprocessing of scRNA-seq, spatial transcriptomics, amyloid beta imaging, and HCR data

We collected four Alzheimer’s Slide-seq mouse hippocampus sections [38] using the Slide-seqV2 protocol [2] (see *Supplementary Methods* for details) on a female 8.8 month old J20 Alzheimer’s mouse model [38]. We used three total Slide-seq mouse cerebellum sections, two collected using the Slide-seqV2 protocol, and one section used from a previous study [24]. Recall that data from multiple sections is integrated as described in *Multiple replicates*. The Slide-seq mouse testes and mouse cancer datasets were used from recent previous studies [12, 49]. In particular, the tumor dataset represented a single *Kras*^G12D/+^ *Trp53*^-/-^ (KP) mouse metastatic lung adenocarcinoma tumor deposit in the liver [50]. The MERFISH hypothalamus dataset was obtained from a publicly available study [11]. To identify cell types on these datasets, we utilized publicly available single-cell RNA-seq datasets for the testes [72], hypothalamus [11], cerebellum [32], cancer [49], and Alzheimer’s hippocampus datasets [73]. All these scRNA-seq datasets have previously been annotated by cell type.

Slide-seq data was preprocessed using the Slide-seq tools pipeline [2]. For all spatial transcriptomics datasets, the region of interest (ROI) was cropped prior to running C-SIDE, and spatial transcriptomic spots were filtered to have a minimum of 100 UMIs. We used prior anatomical knowledge to crop the ROI from an image of the total UMI counts per pixel across space, which in many cases allows one to observe overall anatomical features. For example, in Slide-seq Alzheimer’s hippocampus, the somatosensory cortex was cropped out prior to analysis.

For the Alzheimer’s dataset, in order to test for differential expression with respect to amyloid plaques, we collected fluorescent images of DAPI and amyloid beta (A*β*), using IBL America Amyloid Beta (N) (82E1) A*β* Anti-Human Mouse IgG MoAb on sections adjacent to the Slide-seq data. We co-registered the DAPI image to the adjacent Slide-seq total UMI image using the `ManualAlignImages` function from the `STutility` R package [74]. To calculate plaque density, plaque images were convolved with an exponentially-decaying isotropic filter, using a threshold at the 0. 9 quantile, and normalized to be between 0 and 1. For each Slide-seq section, plaque density was defined as the average between the plaque densities on the two adjacent amyloid sections.

For *in situ* RNA hybridization validation of cerebellum DE results, we collected hybridization chain reaction (HCR) data on genes *Aldoc*, *Kcnd2*, *Mybpc1*, *Plcb4*, and *Tmem132c* (Supplementary Table 9) using a previously developed protocol [75]. We simultaneously collected cell type marker genes of Bergmann (*Gdf10*), granule (*Gabra6*), and Purkinje (*Calb1*) cell types, markers that were sourced from a prior cerebellum study [32]. Data from *Kcnd2* was removed due to the HCR fluorescent channel failing to localize RNA molecules, but rather reflecting tissue autofluorescence. ROIs of nodular and anterior regions were cropped, and background, defined as median signal, was subtracted. For this data, DE was calculated as the log-fold-change, across ROIs, of average gene signal over the pixels within the ROI containing cell type markers of a particular cell type. Pixels containing marker genes of multiple cell types were removed. C-SIDE single-sample standard errors in Figure 3d were calculated by modeling single-sample variance as the sum of the variance across samples and variance representing uncertainty around the population mean.

### Cell type proportion estimation and construction of covariates

For each dataset, we constructed at least one covariate, an axis along which to test for DE. All covariates were scaled linearly to have minimum 0 and maximum 1. For the cerebellum dataset, the covariate was defined as an indicator variable representing membership within the nodular region (as opposed to the anterior region). The nodular and anterior ROIs were annotated manually from the total UMI image, and all other regions were removed. For the testes dataset, the covariate was a discrete variable representing the cellular microenvironment of tubule stage, labels that were obtained from tubule-level gene expression clustering from the previous Slide-seq testes study [12]. In that study and here, tubules are categorized into 4 main stages according to tubule sub-stage groups of stage I–III, IV–VI, VII–VIII, and IX–XII. For the cancer dataset, the covariate was chosen to be the density of the myeloid cell type. Cell type density was calculated by convolving the cell type locations, weighted by UMI number, with an exponential filter. For this dataset, we also ran C-SIDE nonparametrically. For the Alzheimer’s hippocampus dataset, the covariate was chosen to be the plaque density, defined in Section *Collection and preprocessing*. For the MERFISH hypothalamus dataset, the covariate was defined as distance to the midline, and we also considered quadratic functions of midline distance by adding squared distance as an covariate. For the quadratic MERFISH C-SIDE model, we conducted hypothesis testing on the quadratic coefficient. To estimate platform effects and cell type proportions, RCTD was run on *full mode* for the testes dataset, and was run on *doublet mode* for all other datasets with default parameters [24].

### Validation with simulated gene expression dataset

We created a ground truth DE simulation to test C-SIDE on the challenging situation of mixtures between two cell type layers. We tested C-SIDE on a dataset of cell type mixtures simulated from the cerebellum single-nucleus RNA-seq dataset, which was also used as the reference for cell type mapping. We restricted to Purkinje and Bergmann cell types, which are known to spatially colocalize.

In order to simulate a cell type mixture of cell types A (Purkinje) and B (Bergmann), we randomly chose a cell from each cell type, and sampled a predefined number of UMIs from each cell (total 1, 000). We defined two discrete spatial regions (Figure 1a), populated with A/B cell type mixtures. We varied the mean cell type proportion difference across the two regions and also simulated the case of cell type proportions evenly distributed across the two regions. Cell type-specific spatial differential gene expression also was simulated across the two regions. To simulate cell type-specific differential expression in the gene expression step of the simulation, we multiplicatively scaled the expected gene counts within each cell of each cell type. An indicator variable for the two spatial bins was used as the C-SIDE covariate.

### Additional computational analysis

For confidence intervals on data points or groups of data points (Figure 4d, Figure 4g), we used the predicted variance of data points from C-SIDE (see *Supplementary Methods* for details). Likewise, for such analysis we used predicted counts from C-SIDE at each pixel (*Supplementary Methods*). For the testes dataset, a cell type was considered to be present on a bead if the proportion of that cell type was at least 0.25 (Figure 4d). Additionally, cell type and stage-specific marker genes were defined as genes that had a fold-change of at least 1.5 within the cell type of interest compared to each other cell type. We also required significant cell type-specific differential expression between the stage of interest with all other stages (fold-change of at least 1.5, significance at the level of 0.001, Monte Carlo test on Z-scores). Cyclic genes were defined as genes whose minimum expression within a cell type occurred two tubule stages away from its maximum expression, up to log-space error of up to 0.25.

For nonparametric C-SIDE on the tumor dataset, we used hierarchical Ward clustering to cluster quantile-normalized spatial gene expression patterns into 7 clusters. For gene set testing on the tumor dataset, we tested the 50 hallmark gene sets from the MSigDB database [76] for aggregate effects in C-SIDE differential expression estimates for the tumor cell type. For the nonparametric case, we used a binomial test with multiple hypothesis correction to test for enrichment of any of the 7 spatial clusters of C-SIDE-identified significant genes in any of the 50 gene sets. For the parametric case, we used a permutation test on the average value of C-SIDE *Z* scores for a gene set. That is, we modified an existing gene set enrichment procedure [77] by filtering for genes with a fold-change of at least 1.5 and using a two-sided permutation test rather than assuming normality. In both cases, we filtered to gene sets with at least 5 genes and we used Benjamini-Hochberg procedure across all gene sets to control the false discovery rate at 0.05. The proportion of variance not due to sampling noise (Figure 5b) was calculated by considering the difference between observed variance on normalized counts and the expected variance due to Poisson sampling noise.

We considered and tested several simple alternative methods to C-SIDE, which represent general classes of approaches. First, we considered a two-sample Z-test on single cells (defined as pixels with cell type proportion at least 0.9). Additionally, we tested *Bulk* differential expression, which estimated differential expression as the log-ratio of average normalized gene expression across two regions. The *Single* method of differential expression rounded cell type mixtures to the nearest single cell type and computed the log-ratio of gene expression of cells in that cell type. Finally, the *Decompose* method of differential expression used a previously-developed method to compute expected gene expression counts for each cell type [24], followed by computing the ratio of cell type-specific gene expression in each region.

### Implementation details

C-SIDE is publicly available as part of the R package https://github.com/dmcable/spacexr. The quadratic program that arises in the C-SIDE optimization algorithm is solved using the `quadprog` package in R [78]. Prior to conducting analysis on C-SIDE output, all ribosomal proteins and mitochondrial genes were filtered out. Additional parameters used for running C-SIDE are shown in Supplementary Table 10. C-SIDE was tested on a Macintosh laptop computer with a 2.4 GHz Intel Core i9 processor and 32GB of memory (we recommend at least 4GB of memory to run C-SIDE). For example, we timed C-SIDE with four cores on one of the Slide-seq cerebellum replicates, containing 2, 776 pixels across two regions, 5 cell types, and 4, 812 genes. Under these conditions, C-SIDE ran in 13 minutes and 47 seconds (excluding the cell type assignment step in which computational efficiency has been described previously [24]).

## Author Contributions

D.M.C., R.A.I, and F.C. conceived the study; F.C., E.M., E.Z.M., and D.C. designed the Slide-seq, antibody stain, and HCR experiments; E.M. generated the Slide-seq, antibody stain, and HCR data; D. M.C., R.A.I., and F.C. developed the statistical methods; D.M.C., F.C., and R.A.I designed the analysis; D.M.C., S.Z., M.D., R.A.I., and F.C. analyzed the data; D.M.C., F.C, R.A.I., V.S., and H.C. interpreted biological results; V.S. annotated the tumor H&E stain; D.M.C., F.C., and R.A.I. wrote the manuscript; all authors read and approved the final manuscript.

## Acknowledgements

We thank Luli Zou and Robert Stickels for providing valuable input on the analysis. We thank Tongtong Zhao and Zachary Chiang for generously providing the cancer Slide-seq data and providing helpful feedback. We thank Samuel Marsh for kindly providing mouse J20 Alzheimer’s model samples. We thank members of the Chen lab, Irizarry lab, and Macosko lab including Tushar Kamath for helpful discussions and feedback. D.C. was supported by a Fannie and John Hertz Foundation Fellowship and an NSF Graduate Research Fellowship. This work was supported by an NIH Early Independence Award (DP5, 1DP5OD024583 to F.C.), the NHGRI (R01, R01HG010647 to F.C. and E.Z.M), as well as the Burroughs Wellcome Fund, the Searle Scholars Award, and the Merkin Institute to F.C. R.A.I. was supported by NIH grants R35GM131802 and R01HG005220.

## Conflict of Interest Statement

E. Z.M. and F.C. are listed as inventors on a patent application related to Slide-seq. F.C. is a paid consultant for Celsius Therapeutics and Atlas Bio.

## Data Availability Statement

Slide-seq V2 data generated for this study is available at the Broad Institute Single Cell Portal https://singlecell.broadinstitute.org/single_cell/study/SCP1663. Additional publicly available data from other studies that was used for analysis is also included in this repository.

## Code Availability Statement

C-SIDE is implemented in the open-source R package *spacexr*, with source code freely available at https://github.com/dmcable/spacexr. Additional code used for analysis in this paper is available at https://github.com/dmcable/spacexr/tree/master/AnalysisC-SIDE.

## Supplementary Methods

### Introduction and model definition

We now revisit our Cell type-Specific Inference of Differential Expression (C-SIDE) model at an increased level of detail. Recall the following definition of the C-SIDE model, where for each pixel *i* = 1,…, *I* in the spatial transcriptomics dataset, we denote the observed gene expression counts as *Y_i,i,g_* for each gene *j* = 1,…, *J* and experimental sample *g* = 1,…, *G*:

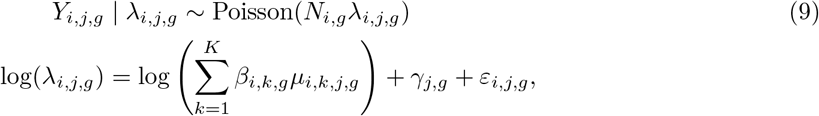

with *N_i,g_* the total transcript count or number of unique molecular identifies (UMIs) for pixel *i* and sample *g*, *K* the number of cell types present in our dataset, *μ_k,j,g_* the mean gene expression profile for cell type *k* and gene *j* and sample *g*, *β_i,k,g_* the proportion of the contribution of cell type *k* to pixel *i* in sample *g*, *γ_j,g_* a gene-specific platform random effect, and *ε_i,j,g_* a random effect to account for other technical and biological sources of variation. We assume *γ_j,g_* and *ε_i,j,g_* both follow normal distributions with mean 0 and standard deviation *σ_γ,g_* and *σ_ε,j,g_*, respectively. Lastly, *μ_i,k,j,g_* represents the average gene expression of gene *j* in cell type *k* at pixel location *i* in sample *g*. We model *μ_i,k,j,g_*, for each gene *j*, each cell type *k*, and each sample *g* as depending log-linearly on several covariates, *x*:

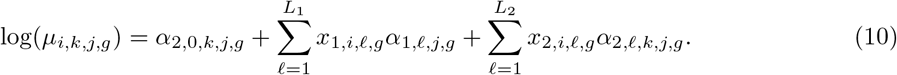

More specifically, we split our covariates into two sets (of sizes *L*_1_ and *L*_2_). The first set, *x*_1,*i,ℓ,g*_, share coefficients across cell types, while the second set, *x*_2,*i,ℓ,g*_, has a different coefficient for each cell type. This notation is different from the presentation of C-SIDE in the main methods section, in which *x*_1_ was not present and no coefficients were shared across cell types. In practice, we do not typically assume that differential expression is shared across cell types (that is, *x*_1,*i,ℓ,g*_ is not used), but *x*_1_ is included here as an optional feature. We have *x*.,_*i,ℓ,g*_ representing the *ℓ*’th covariate, evaluated at pixel *i* in sample *g*. In all cases, *x* is pre-determined to contain variables on which gene expression is hypothesized to depend.

For each covariate *x*, there is a corresponding coefficient *α*. More precisely, *α*_1,*ℓ,j,g*_ represents a gene expression change per unit change of *x*_1,*i,ℓ,g*_ for gene *j* in sample *g*. Note that this coefficient is the same across all cell types. On the other hand, *α*_2,*ℓ,k,j,g*_ represents a gene expression change per unit change of *x*_2,*i,ℓ,g*_ specific to cell type *k* in sample *g*. Finally, *α*_2,0,*k,j,g*_ represents the intercept term for gene *j* and cell type *k* in sample *g*. For ease of notation, we will sometimes use *α*_1,*ℓ,k,j,g*_ to equal *α*_1,*ℓ,j,g*_ for all *k*. Moreover, we will use *α* to refer to the joint vector of both *α*_1_ and *α*_2_. The parameters *α* are estimated by C-SIDE by maximum likelihood. C-SIDE also obtains standard errors for each coefficient *α*. These standard errors are subsequently used for confidence intervals and hypothesis testing.

### Maximum Likelihood Estimation

C-SIDE estimates the parameters of (9) via maximum likelihood estimation. First, we note that all parameters in the model are independent across samples. As such, we fit the model independently for each sample, and we now drop the subscript of sample *g* for notational convenience. We will return to the issue of integrating results across multiple samples in Multiple replicates. First, the parameters *β_i,k_* and *γ_j_* are estimated by the RCTD algorithm as previously described [24]. We can accurately estimate cell type proportions and platform effects without being aware of differential spatial gene expression because differential spatial gene expression is smaller than gene expression differences across cell types. After identifying cell types, C-SIDE estimates gene-specific overdispersion *σ_ε,j_* for each gene by maximum likelihood estimation (see *Fitting the overdispersion parameter*). Finally, C-SIDE estimates the parameters *α*_1,*ℓ,j*_ and *α*_2,*ℓ,k,j*_ by maximum likelihood estimation. For the final key step of estimating *α*, we use plugin estimates (denoted by ^) of *β_i,k_*, *γ_j_*, and *σ_ε_*. After we substitute (10) into (9), we obtain:

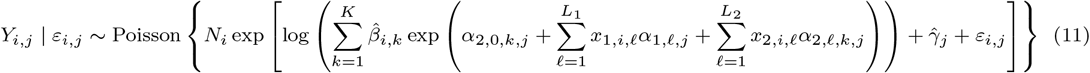

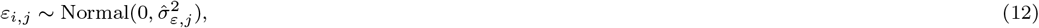

Now, we provide an algorithm for computing the maximum likelihood estimator of *α*. Our likelihood optimization algorithm is a second-order, trust-region based optimization. In brief, we iteratively solve quadratic approximations of the log-likelihood, adaptively constraining the maximum parameter change at each step. Critically, the likelihood is independent for each gene, so separate genes can be run in parallel.

Now, we consider the computation of the maximum likelihood estimator (MLE) of *α* for the likeli-hood 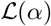 of observing *Y_i_* for 1 ≤ *i* ≤ *I*, using the assumption that measurements on separate pixels are independent. We define the predicted counts at pixel *i* as 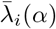, where,

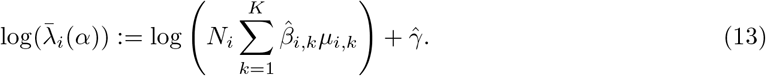

From now on, we will drop the constant term 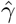, as it can be equivalently factored into the *μ* intercept term. Next, we can use (9) to compute the likelihood of the C-SIDE model,

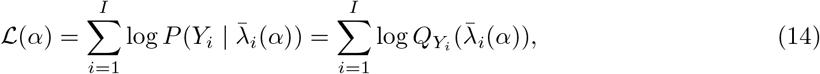

where we have introduced the function *Q* to represent the probability, under our Poisson-log-normal sampling model, of observing *Y_i_* counts given predicted counts *λ_i_*(*α*),

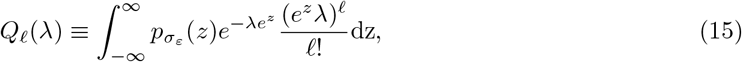

where *p_σ_ε__* is the normal distribution pdf with standard deviation *σ_#x03B5;_*. To optimize our likelihood, we develop a second-order trust-region optimization method [79], in which sequential quadratic approximations are optimized within a trust region, whose size is determined adaptively. To do so, we first initialize *α* as *α*_0_, which is set to 0 for intercept terms, and –5 for non-intercept terms. Additionally, we initialize the trust-region width, *δ*, as *δ*_0_ = 0.1. At step *n* + 1 of the algorithm, with previous parameters *α_n_* and *δ_n_*, we make the following quadratic Taylor approximation, 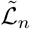 to 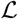,

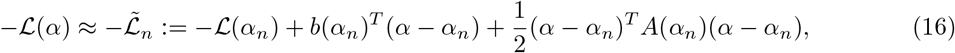

where *b* and *A* represent the gradient and Hessian of 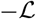, respectively, which are computed below. Next, we define 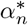 as the solution to the following optimization problem of this quadratic approximation over the trust region:

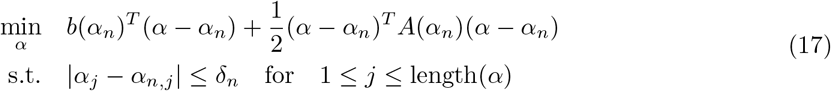

This quadratic program is solved using the `quadprog` package in R [78]. Next, we define *α*_*n*+1_ as:

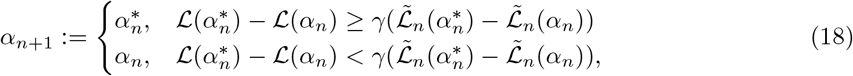

where *γ* = 0.1. Additionally, the trust region is updated as:

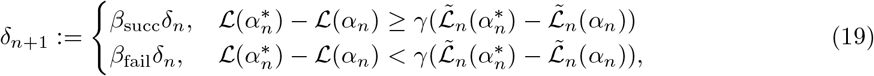

where *β*_succ_ = 1.1 and *β*_fail_ = 0.5, which, along with *γ*, were chosen by a combination of using standard parameter choices [79] and ensuring efficient and stable convergence to local minima. Intuitively, the quadratic approximation 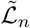 will only be accurate within a local region, and the trust region is intended to empirically approximate that region. In order to test whether our local approximation is accurate, we check whether the predicted gain in log-likelihood, 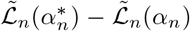, is close to the true gain in log-likelihood, 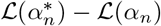, within a factor of *γ*. If the local approximation is indeed accurate, the algorithm takes a step, and the trust region is allowed to grow. If not, the algorithm stays put, and the trust region shrinks. This prevents the algorithm from diverging due to poor quadratic approximations. This procedure is repeated until convergence (see *Stopping conditions and convergence*).

### Gradient and Hessian

In this section, we will derive an expression for the gradient and hessian of 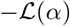. First, we can calculate the gradient as,

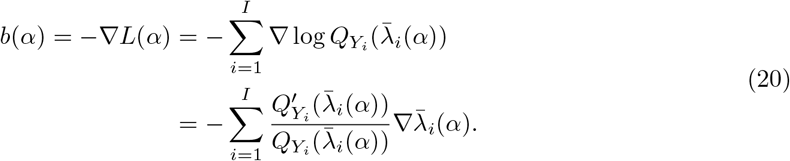

Additionally, we have the Hessian,

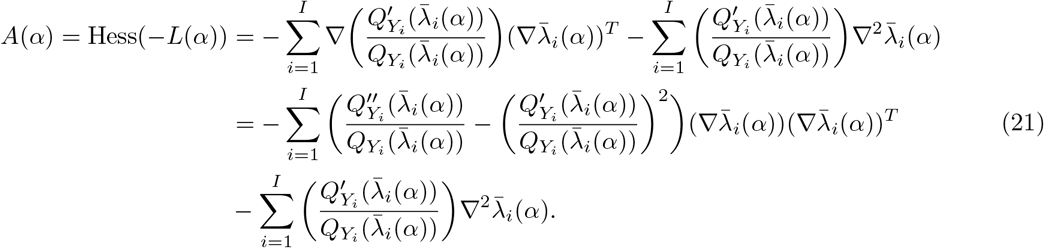

We recall the procedure for computing *Q* and its derivatives as previously described [24]. What remains is to calculate explicit expressions for 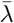 and its derivatives, which we do now. From (10) and (15), we recall the definition of 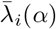:

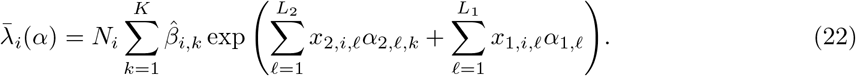

Next, we calculate the gradient of 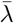 with respect to *α*_1_ and *α*_2_ separately:

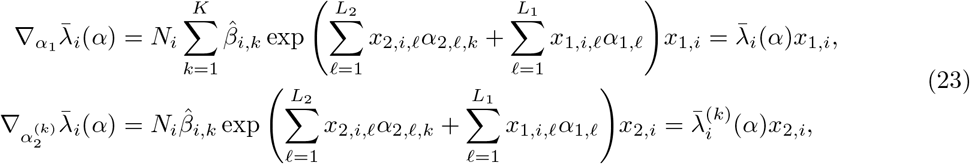

where we have defined 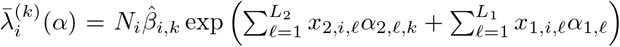. Next, we can compute the second derivatives:

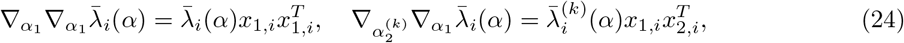

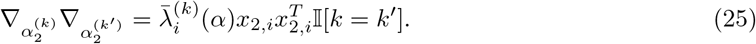

Finally, notice that all the above expressions, including 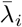 and 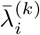 across all pixels *i*, can be computed efficiently using matrix multiplications. Lastly, the Fisher information is computed as a scaled version of the Hessian (see *Justification of consistency and asymptotic normality*).

### Stopping conditions and convergence

The algorithm stops when one of two conditions are satisfied: *δ_n_* < *ε*_1_ or 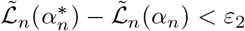 for 6 consecutive iterations. Default choices are *ε*_1_ = .001 and *ε*_2_ = .00001. Assume that the algorithm stops after *n*–1 iterations and arrives at solution *α_n_*. Convergence is defined by considering the distance of *α_n_* to the optimal solution of 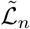, which is the maximum step size of the next step of the algorithm. Since 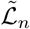 is a quadratic function, its maximum can be calculated as *α*\*:= *α_n_* – *A*(*α_n_*)^-1^*b*(*α_n_*). Consequently, *α_n_* – *α*\* = *A*(*α_n_*)^-1^*b*(*α_n_*). For each parameter 1 ≤ *i* ≤ length(*α*), we define that parameter *i* has converged if 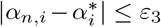, where *ε_3_* = .01. Intuitively, for all parameters *i* such that 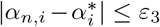, these parameters will change by at most *ε_3_* in the next step of the algorithm. Note that it is possible for some parameters to converge while others do not. In the most common scenario, consider a case in which one cell type has very low gene expression in the gene of interest. In this case, it is possible that the parameter controlling the expression of this gene will diverge to –∞. As such, this parameter doesn’t have a practical effect on the model, but it should not prevent the other parameters (of cell types with higher expression) from converging. For each cell type, we filter out genes that did not converge for downstream analysis. In the multi-region case, for each cell type, we test for differential expression among the subset of regions that have converged.

### Fitting the overdispersion parameter

Here, we describe the procedure for fitting the gene-dependent overdispersion parameter *σ_ε,j_*. This is necessary because we found evidence that the overdispersion depends on gene *j*, and modeling genespecific overdispersion is necessary for controlling the false-positive rate of C-SIDE. In order to fit a gene-dependent overdispersion parameter, we fit C-SIDE with initial overdispersion parameter *σ_ε_*, which is obtained from the cell type identification step. Next, we use the fitted parameters *α* and calculate the log-likelihood of C-SIDE for each possible choice of *σ* (out of a discrete set ranging from 0.1 to 2). Because the log-normal distribution has a mean of *e*^*σ*^2^/2^, the C-SIDE predicted expression values 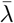 are scaled by *e*^-*σ*^2^/2^ to maintain a consistent mean across different values of *σ*. In practice, this decision substantially increases the rate of convergence. After computing log-likelihood values for each *σ*, the best *σ* is chosen, and the parameters of C-SIDE are re-fit. This procedure is repeated until convergence at *σ* = *σ_ε,j_*.

### Predicted mean and variance of individual data pixel counts

After *α* is estimated, we can compute the predicted mean and variance of *Y_i_*, given *x_i_*, according to the C-SIDE model. These predictions are used to check whether the observed behavior of data points agrees with the predictions of the C-SIDE model. Rewriting (9),

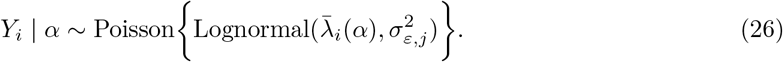

Using properties of the lognormal distribution, we can calculate the mean counts,

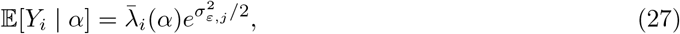

as well as the variance of the counts, using the law of total variance,

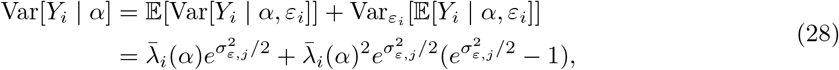

where the first part used the equivalence of the mean and variance of the Poisson distribution, and the second part used the variance of the lognormal distribution.

### Multiple replicates

In order to extend the hypothesis testing framework to the case of multiple replicates, we now recall *α_g_* and *s_g_* to be the differential expression and standard error for replicate *g*, where 1 ≤ *g* ≤ *G*, and *G* > 1 is the total number of replicates. We will consider testing for differential expression for fixed covariate *ℓ*, cell type *k*, and gene *j*. In this case, as later derived in (53), the observed estimate 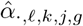, conditional on *α*, follows a univariate normal distribution with standard deviation *s._ℓ,k,j,g_*:

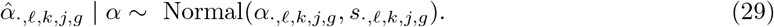

We further assume that additional biological and/or technical variation across samples exists, such that each *α_g_* is normally distributed around a population-level differential expression *A*, with standard deviation *τ*:

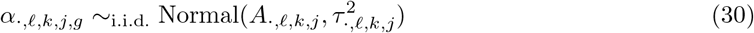

We estimate *τ* using the method of moments (second moment) on the observed estimate 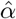, obtained independently from each sample:

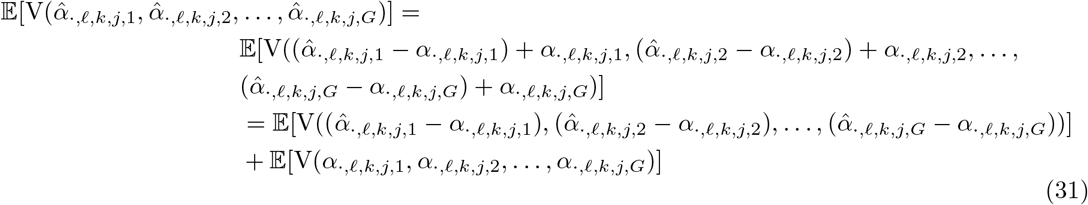

Here, the second step utilizes the independence of 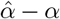 and *α*. Additionally, we use the finite sample variance function V to denote 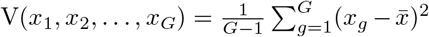, which is an unbiased estimator of the variance of *x* if *x_g_* is an i.i.d. random variable. Consequently, the second term above equals 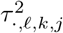. Additionally, since 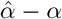 is mean 0, we can use the fact that for mean 0 variables *y* that are coordinate-wise independent, 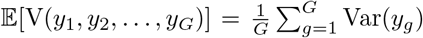. Applying this fact to the first term, we obtain,

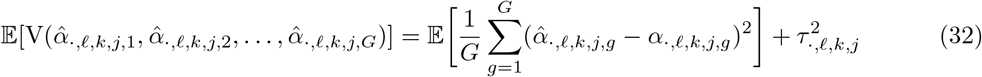

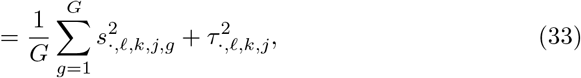

where we have used the C-SIDE standard errors *s*^2^ to estimate the variance of 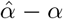. Consequently, we obtain the following method of moments estimator of *τ*^2^:

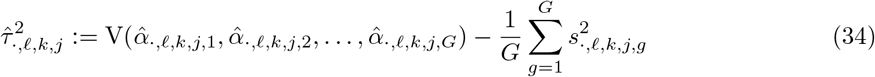

Given the above analysis, the estimator is the unbiased method of moments estimator. Since we know that *τ*^2^ is nonnegative, we next modify our estimator to an estimator that dominates the original:

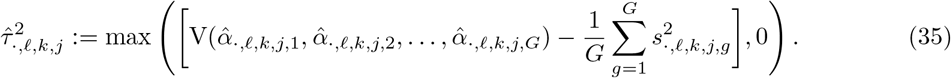

We note that the above method of moments estimator (and our overall approach) is similar to the widely used DerSimonian-Laird method in meta-analysis [70, 71]. After utilizing the estimate of *τ*^2^, we can now compute the estimate and standard error of *A*, as follows. Given equations, (29) and (30), we have that 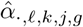 is distributed independently for 1 ≤ *g* ≤ *G* as:

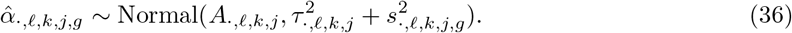

By the Gauss-Markov theorem for Generalized Least Squares, the best (i.e. minimum variance) unbiased estimator of *A* is:

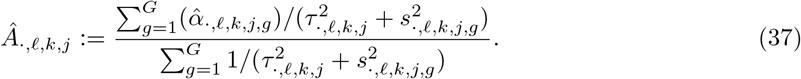

We further plugin our estimate 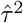 for *τ*^2^, which is an approach called feasible generalized least squares:

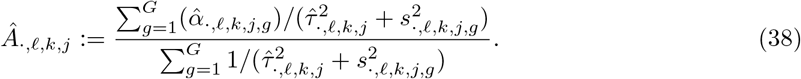

Finally, the feasible estimate of variance of this estimator (also by the Gauss-Markov theorem) is:

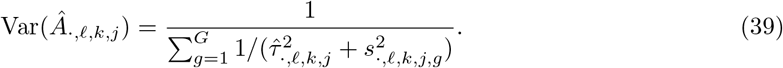

### Multiple samples and replicates

After developing a hypothesis testing framework for the case of multiple replicates, we now consider the extension of this framework to the more complicated study design of multiple biological samples (*M* samples) with multiple replicates per sample (*G_m_* replicates per sample). In this case, we now model *α* for each sample 1 ≤ *m* ≤ *M* and each replicate 1 ≤ *g* ≤ *G_m_* as normally distributed, independently for each replicate, with standard deviation *τ*, as follows,

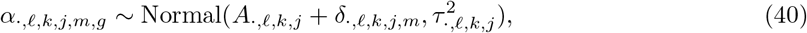

where *δ* represents a sample-specific random effect which is itself normally distributed with standard deviation Δ,

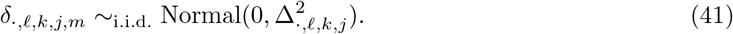

Notice that for fixed sample *m*, conditional on *δ*, our problem is identical to the multiple replicate case above, given a population-mean of *A*._*ℓ,k,j*_ + *δ*._*ℓ,k,j,m*_. Using this reasoning, we take as an estimate of *τ*^2^ the average, across samples, of the estimates of *τ*^2^ in (35). As we have utilized the variance within each sample to obtain an estimate of *τ*, we will next use the variance across samples to estimate *Δ*. We take (38) and (39) as the value and variance (conditional on *δ*) respectively of the following unbiased estimate *E* of *A*._*ℓ,k,j*_ + *δ*.,_*ℓ,k,j,m*_, which represents the differential expression within sample *m*,

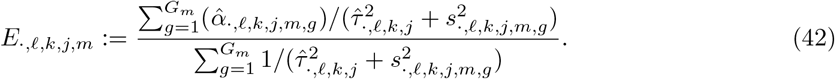

Given that *E._ℓ,k,j,m_* is an unbiased estimate of *A.,_ℓ,k,j_* + *δ.,_ℓ,k,j,m_*, we recognize that our problem has been reduced to the original multiple replicates problem (addressed above), where *α* has been replaced with *A* + *δ*, *τ* has been replaced with Δ, 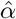 has been replaced by *E*, and *s*^2^ has been replaced by what we define as *S*^2^, the conditional (on *δ*) variance of *E* given in (39),

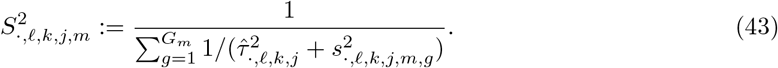

As a result of this observation, we can apply a similar derivation as that of (35) to obtain the following method of moments estimate of Δ,

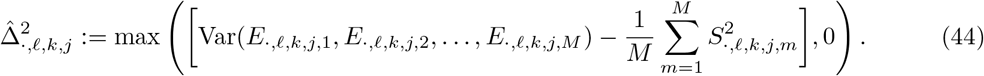

Continuing our parallel to our previous result, we use the feasible Gauss-Markov estimator of *A* derived in in (38) and (39),

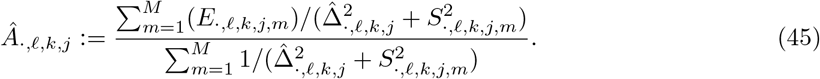

Moreover, using (39), the feasible estimate of variance of this estimator is,

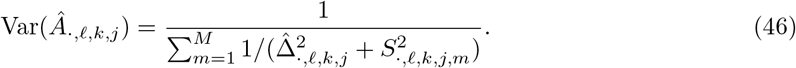

Therefore, we have derived estimators of population-level differential expression in the case of multiple eplicates or multiple samples with multiple replicates.

### Justification of consistency and asymptotic normality of maximum likelihood estimator of *α*

Since each gene and each sample analyzed independently, we drop the notation of gene *j* and sample *g*. First, we consider the joint distribution of all the variables in our model: *x_i_*, *β_i_*, and *Y_i_*. We recall hat *x_i_* and *Y_i_* are observed, and we assume that these variables are generated i.i.d. for each pixel 1 ≤ *i* ≤ *n*, with *n*:= *I*):

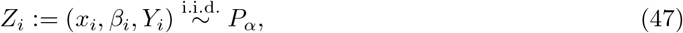

where *Z_i_* represents the joint random variable and *P_α_*(*Z_i_*) = *Q*(*x_i_*, *β_i_*)*P_α_*(*Y_i_*| *x*, *β*). Here, *Q* represents the joint distribution, across pixels, of cell type proportions and covariates, which we assume does not depend on *α*. As estimation of *α* does not depend on this term, we will ignore this term. The conditional distribution *P_α_*(*Y_i_*| *x*, *β*) is precisely the probabilistic model specified by C-SIDE in (9). For this analysis, we treat *β* as observed and do not consider the uncertainty around the estimation of *β*, as errors in the estimation of *β* are expected to be small and independent across pixels.

Due to the specification of C-SIDE, assuming that the columns of *x* are linearly independent, identifiability is satisfied. That is, *P_α_* ≠ *P_α′_* for any other pair of distinct parameters *α* and *α*′. It follows from standard asymptotic theory results [80] (using additional regularity conditions including Lipschitz continuity of second derivatives and local convexity of the C-SIDE log-likelihood within a bounded region) that if we let 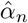 be the MLE estimator on *n* pixels, then asymptotic consistency holds:

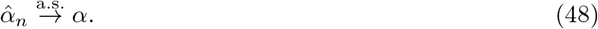

In addition to consistency, asymptotic normality holds as *n* → ∞ [80]:

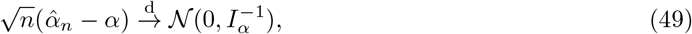

where *I_α_* is defined to be the Fisher information, which can be represented as,

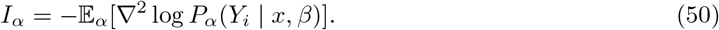

In our case, we will use the observed Fisher information *Î_α_* to estimate the Fisher information:

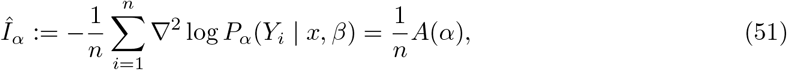

where *A*(*α*), defined in (16), is the Hessian of the C-SIDE log-likelihood function. Substituting the Hessian into the equation (49) above, we conclude that approximately for large *n*,

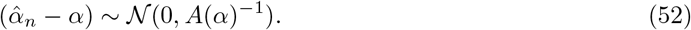

Next, for a fixed individual cell type *k*, gene *j*, sample *g*, and covariate *ℓ*, the distribution of 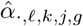 follows a univariate normal distribution with standard deviation *s.,_ℓ,k,j,g_*. According to (49), if we define *s* as 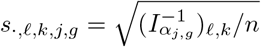, we conclude that,

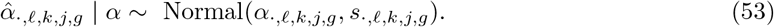

Thus, we have derived the asymptotic distribution of 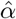, allowing us to compute confidence intervals and perform statistical inference.

## Supplementary Experimental Methods

### Animal Handling

All procedures involving animals at the Broad Institute were conducted in accordance with the US National Institutes of Health Guide for the Care and Use of Laboratory Animals under protocol number 0120-09-16.

### Transcardial Perfusion

C57BL/6J mice were anesthetized by administration of isoflurane in a gas chamber flowing 3% isoflurane for 1 minute. Anesthesia was confirmed by checking for a negative tail pinch response. Animals were moved to a dissection tray and anesthesia was prolonged via a nose cone flowing 3% isoflurane for the duration of the procedure. Transcardial perfusions were performed with ice cold pH 7.4 HEPES buffer containing 110 mM NaCl, 10 mM HEPES, 25 mM glucose, 75 mM sucrose, 7.5 mM MgCl2, and 2.5 mM KCl to remove blood from brain and other organs sampled. The appropriate organs were removed and frozen for 3 minutes in liquid nitrogen vapor and moved to −80C for long term storage.

### Tissue Handling

Fresh frozen tissue was warmed to −20 C in a cryostat (Leica CM3050S) for 20 minutes prior to handling. Tissue was then mounted onto a cutting block with OCT and sliced at a 5° cutting angle at 10 μm thickness. Pucks were then placed on the cutting stage and tissue was maneuvered onto the pucks. The tissue was then melted onto the puck by moving the puck off the stage and placing a finger on the bottom side of the glass. The puck was then removed from the cryostat and placed into a 1.5 mL eppendorf tube. The sample library was then prepared as below. The remaining tissue was re-deposited at −80 C and stored for processing at a later date.

### Puck preparation and sequencing

Pucks were prepared as described recently using barcoded beads synthesized in-house on an Akta Oligopilot 10 according to the updated Slide-seqV2 protocol [2]. Pucks were sequenced using a monobase-encoding sequencing-by-ligation approach also described in the updated protocol. We used slide-seq tools for alignment and processing of Slide-seq data.

Pucks were generated using one of two separate bead batches with the oligo sequences listed below:

Batch 1:

5’-TTT_PC_GCCGGTAATACGACTCACTATAGGGCTACACGACGCTCTTCCGATCTJJJJJJJJTCTTCAGCGTTCCCGAGAJ JJJJJJTCNNNNNNNNT25

Batch 2:

5’-TTT_PC_GCCGGTAATACGACTCACTATAGGGCTACACGACGCTCTTCCGATCTJJJJJJJJTCTTCAGCGTTCCCGAGAJ JJJJJNNNNNNNVVT30

“PC” designates a photocleavable linker; “J” represents bases generated by split-pool barcoding, such that every oligo on a given bead has the same J bases; “N” represents bases generated by mixing, so every oligo on a given bead has different N bases; and “TX” represents a sequence of *x* thymidines. “V” represents bases which may contain A, C, G but not T.

### Slide-seqV2 library preparation

#### RNA Hybridization

Pucks in 1.5 mL tubes were immersed in 200 μL of hybridization buffer (6x SSC with 2 U/μL Lucigen NxGen RNAse inhibitor) for 15 minutes at room temperature to allow for binding of the RNA to the oligos on the beads.

#### First Strand Synthesis

Subsequently, first strand synthesis was performed by incubating the pucks in RT solution for 30 minutes at room temperature followed by 1.5 hours at 52 °C.

#### RT solution

115 μL H2O

40 μL Maxima 5x RT Buffer (Thermofisher, EP0751)

20 μL 10 mM dNTPs (NEB N0477L)

5 μL RNase Inhibitor (Lucigen 30281)

10 μL 50 μM Template Switch Oligo (Qiagen #339414YCO0076714)

10 μL Maxima H-RTase (Thermofisher, EP0751)

#### Tissue Digestion

200 μL of 2x tissue digestion buffer was then added directly to the RT solution and the mixture was incubated at 37 °C for 30 minutes.

#### 2x tissue digestion buffer

200 mM Tris-Cl pH 8

400 mM NaCl

4% SDS

10 mM EDTA

32 U/mL Proteinase K (NEB P8107S)

#### Second Strand Synthesis

The solution was then pipetted up and down vigorously to remove beads from the surface, and the glass substrate was removed from the tube using forceps and discarded. 200 μL of Wash Buffer was then added to the 400 μL of tissue clearing and RT solution mix and the tube was then centrifuged for 2 minutes at 3000 RCF. The supernatant was then removed from the bead pellet, the beads were resuspended in 200 μL of Wash Buffer, and were centrifuged again. This was repeated a total of three times. The supernatant was then removed from the pellet. The beads were then resuspended in 200 μL of ExoI mix and incubated at 37 °C for 50 minutes.

#### Wash Buffer

10 mM Tris pH 8.0

1 mM EDTA

0.01% Tween-20

#### ExoI mix

170 μL H20

20 μL ExoI buffer

10 μL ExoI (NEB M0568)

After ExoI treatment the beads were centrifuged for 2 minutes at 3000 RCF. The supernatant was then removed from the bead pellet, the beads were resuspended in 200 μL of Wash Buffer, and were centrifuged again. This was repeated a total of three times. The supernatant was then removed from the pellet. The pellet was then resuspended in 200 μL of 0.1 N NaOH and incubated for 5 minutes at room temperature. To quench the reaction, 200 μL of Wash Buffer was added and beads were centrifuged for 2 minutes at 3000 RCF. The supernatant was then removed from the bead pellet, the beads were resuspended in 200 μL of Wash Buffer, and were centrifuged again. This was repeated a total of three times. Second Strand Synthesis was then performed on the beads by incubating the pellet in 200 μL of Second Strand Mix at 37 °C for 1 hour.

#### Second Strand Synthesis mix

133 μL H2O

40 μL Maxima 5x RT Buffer

20 μL 10 mM dNTPs

2 μL 1 mM dN-SMRT oligo

5 μL Klenow Enzyme (NEB M0210)

After Second Strand Synthesis, 200 μL of Wash Buffer was added and the beads were centrifuged for 2 minutes at 3000 RCF. The supernatant was then removed from the bead pellet, the beads were resuspended in 200 μL of Wash Buffer, and were centrifuged again. This was repeated a total of three times.

#### Library Amplification

200 μL of water was then added to the bead pellet and the beads were centrifuged for 2 minutes at 3000 RCF. The supernatant was then removed from the bead pellet and the beads were resuspended in 50 μL of library PCR mix and moved into a 200 μL PCR strip tube. PCR was then performed as outlined below:

#### Library PCR mix

22 μL H2O

25 μL of Terra Direct PCR mix Buffer (Takara Biosciences 639270)

1 μL of Terra Polymerase (Takara Biosciences 639270)

1 μL of 100 μM Truseq PCR primer (IDT)

1 μL of 100 μM SMART PCR primer (IDT)

#### PCR program

95 °C 3 minutes

4 cycles of:

98 °C 20 seconds

65 °C 45 seconds

72 °C 3 minutes

9 cycles of:

98 °C 20 seconds

67 °C 20 seconds

72 °C 3 minutes

Then:

72 °C 5 minutes

Hold at 4 °C

#### PCR cleanup and Nextera Tagmentation

Samples were cleaned with Ampure XP (Beckman Coulter A63880) beads in accordance with manufacturer’s instructions at a 0.6x bead/sample ratio (30 μL of beads to 50 μL of sample) and resuspended in 50 μL of water. The cleanup procedure was repeated, this time resuspending in a final volume of 10 μL. 1 μL of the library was quantified on an Agilent Bioanalyzer High sensitivity DNA chip (Agilent 5067-4626). Then, 600 pg of cDNA was taken from the PCR product and prepared into Illumina sequencing libraries through tagmentation using the Nextera XT kit (Illumina FC-131-1096). Tagmentation was performed according to manufacturer’s instructions and the library was amplified with primers Truseq5 and N700 series barcoded index primers. The PCR program was as follows:

#### PCR program

72 °C for 3 minutes

95 °C for 30 seconds

12 cycles of:

95 °C for 10 seconds

55 °C for 30 seconds

72 °C for 30 seconds

72 °C for 5 minutes

Hold at 4 °C

Samples were cleaned with Ampure XP (Beckman Coulter A63880) beads in accordance with manufacturer’s instructions at a 0.6x bead/sample ratio (30 μL of beads to 50 μL of sample) and resuspended in 10 μL of water. 1 μL of the library was quantified on an Agilent Bioanalyzer High sensitivity DNA chip (Agilent 5067-4626). Finally, the library concentration was normalized to 4 nM for sequencing. Samples were sequenced on the Illumina NovaSeq S2 flowcell 100 cycle kit with 12 samples per run (6 samples per lane) with the read structure 44 bases Read 1, 8 bases i7 index read, 50 bases Read 2. Each puck received approximately 200-400 million reads, corresponding to 3,000-5,000 reads per bead.

## Supplementary Figures

**Supplementary figure 1:**
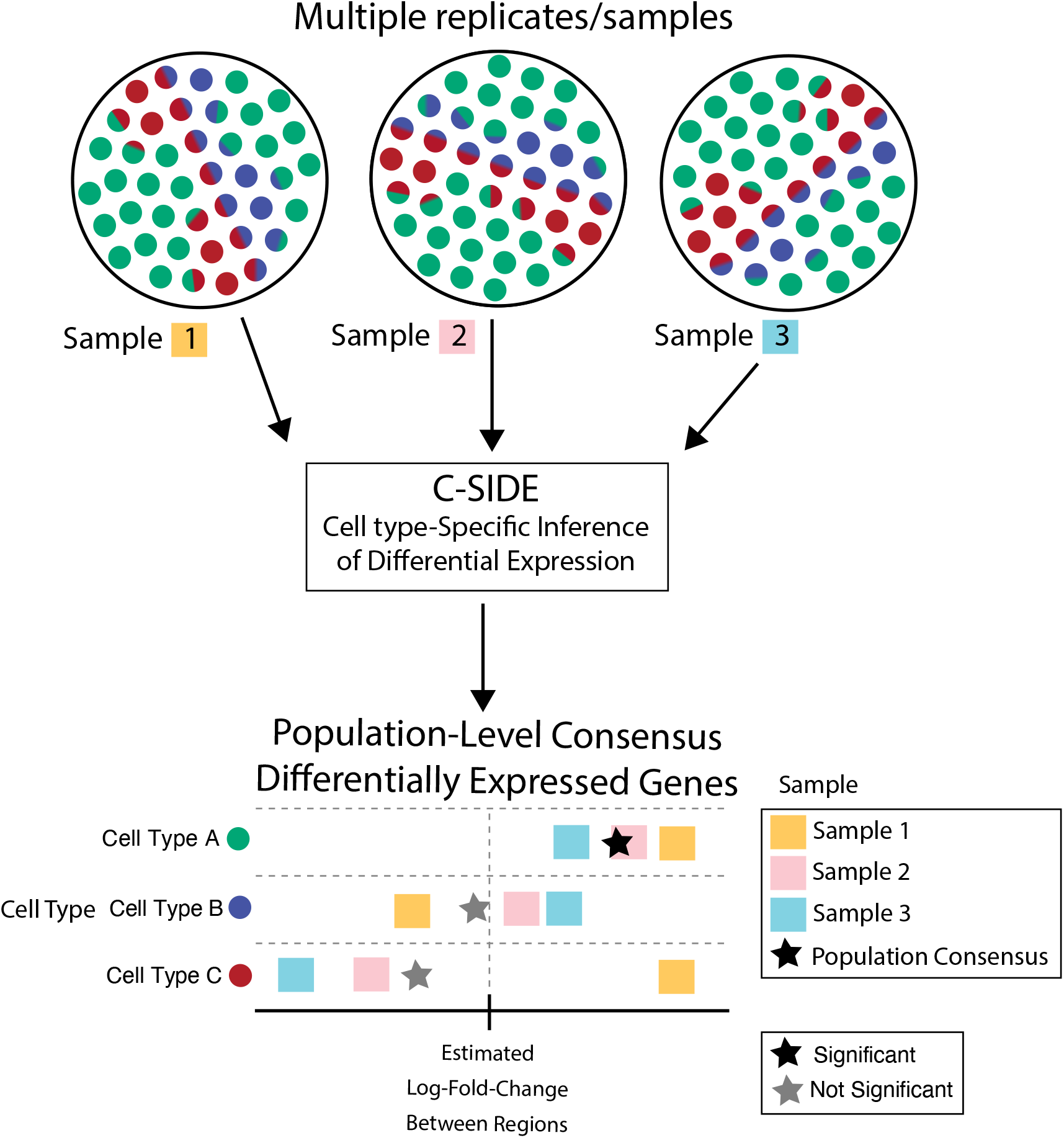
C-SIDE can integrate results from multiple samples to form a robust estimate of population-level consensus differentially-expressed genes.

**Supplementary figure 2:**
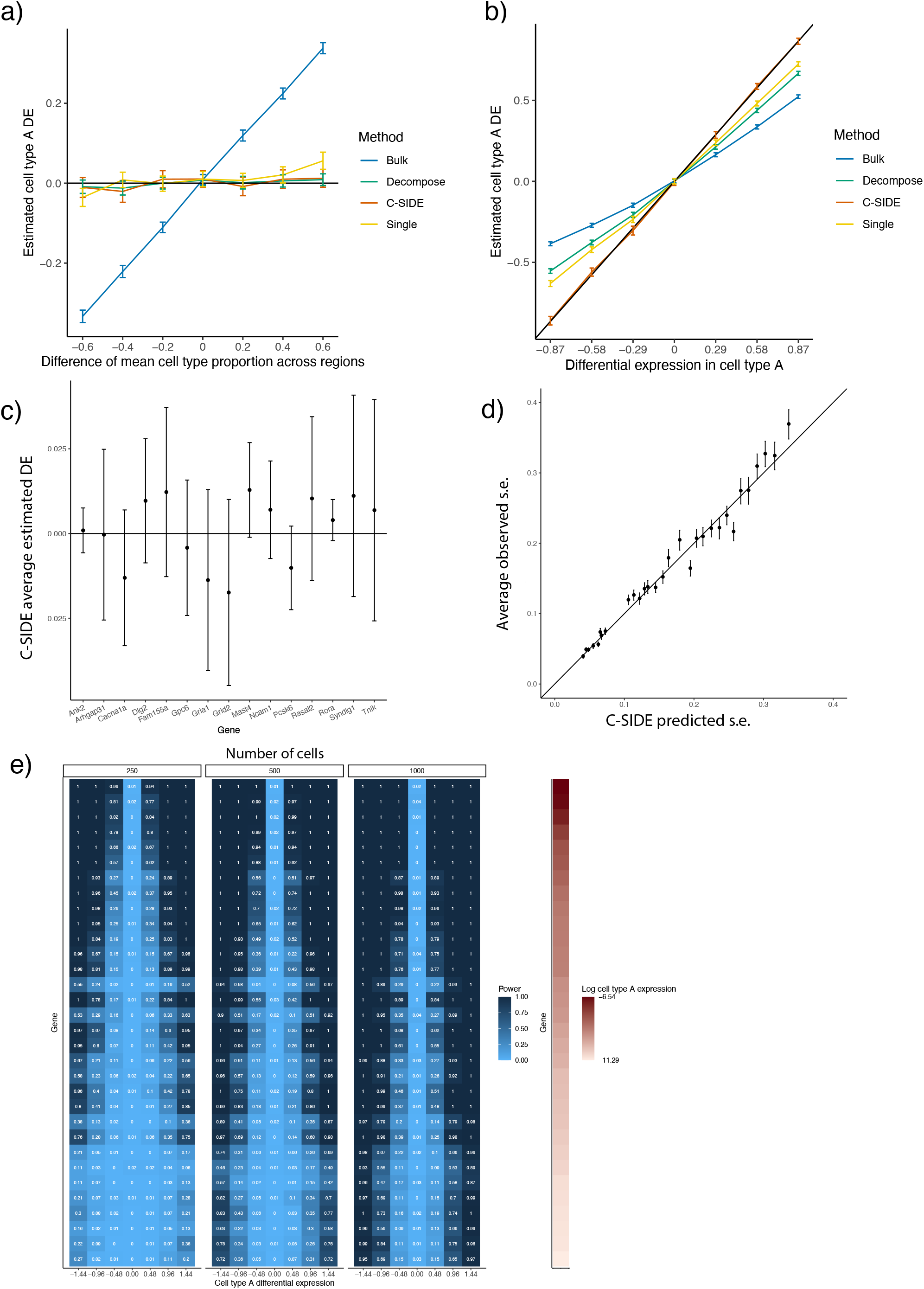
In simulated data, C-SIDE provides unbiased estimates of cell type-specific differential expression, with calibrated *p*–values. All: C-SIDE was tested on a dataset of simulated mixtures of single cells from a single-nucleus RNA-seq cerebellum dataset. (a) Mean estimated cell type A *Astn2* DE (differential expression) across two regions as a function of the difference in mean cell type proportion across regions. Ground truth 0 spatial DE is simulated, and average of (*n* = 100) estimates is shown, along with standard errors. Black line represents ground truth 0 DE (cell type B). Four methods are shown: *Bulk*, *Decompose*, *Single*, and *C-SIDE* (see *Methods* for details). (b) Same as (b) for *Nrxn3* cell type A differential gene expression as a function of DE in cell type A, where *Nrxn3* is simulated to have DE within cell type A but no DE in cell type B. Ground truth identity line shown. (c) C-SIDE mean estimated cell type B differential expression as a function of gene (average over *n* = 500 replicates, with confidence intervals shown). Ground truth line (0 DE) is shown, and each condition used a different gene (out of 15 total genes). (d) Average measured standard error of C-SIDE estimates for each bin of C-SIDE predicted standard error. (e) Statistical power (FPR = 0.01) as a function of gene (y-axis), cell type A DE (x-axis), and number of cells (table number). Genes are sorted by cell type A expression (shown on right in log2 counts per 1).

**Supplementary figure 3:**
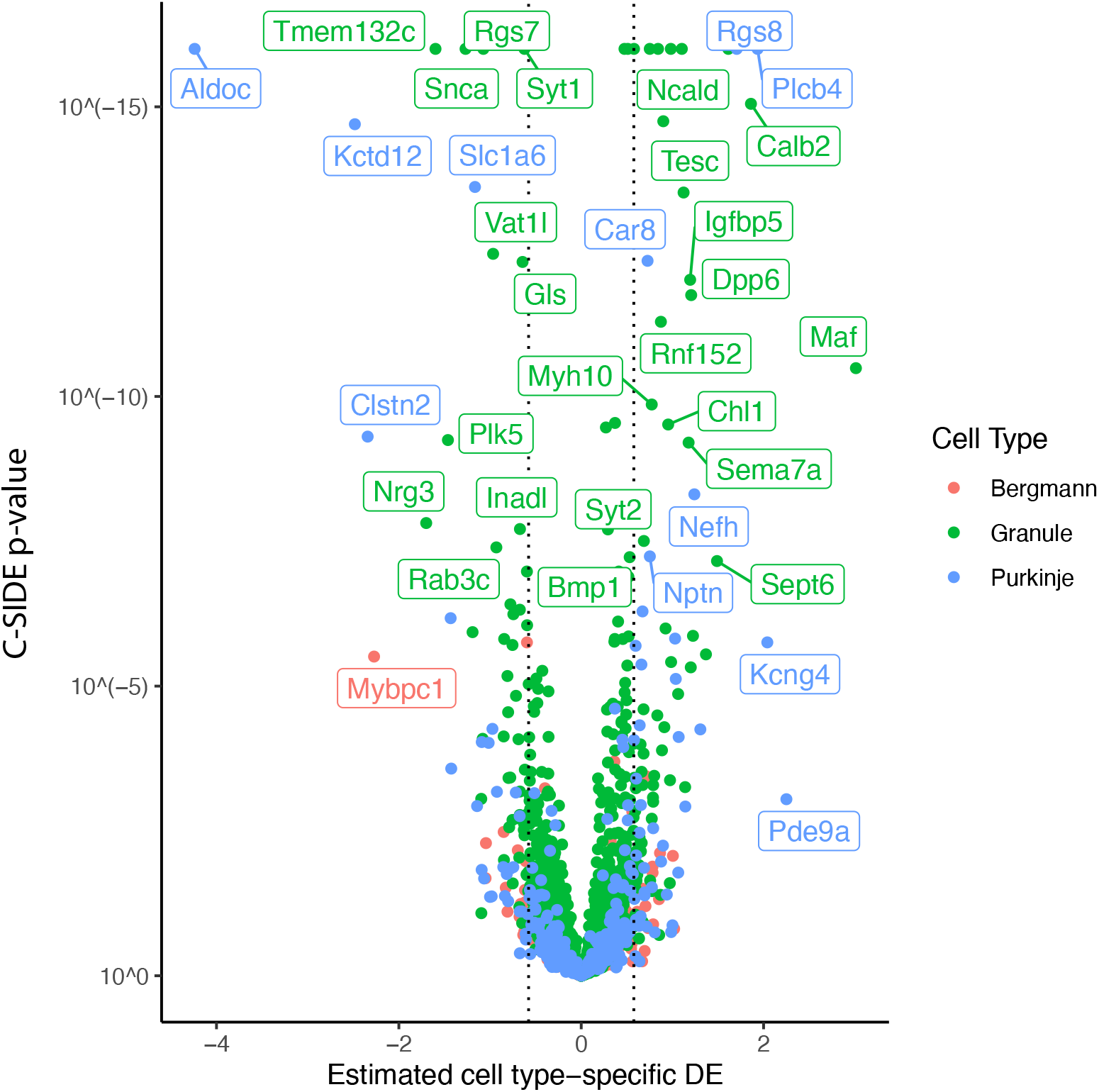
Volcano plot of C-SIDE log2 differential expression results for cerebellum Slide-seq across three replicates, with positive values representing enrichment in the anterior region vs. the nodulus. Color represents cell type, and a subset of significant genes are labeled. Dotted lines represents C-SIDE fold-change cutoff at 1.5.

**Supplementary figure 4:**
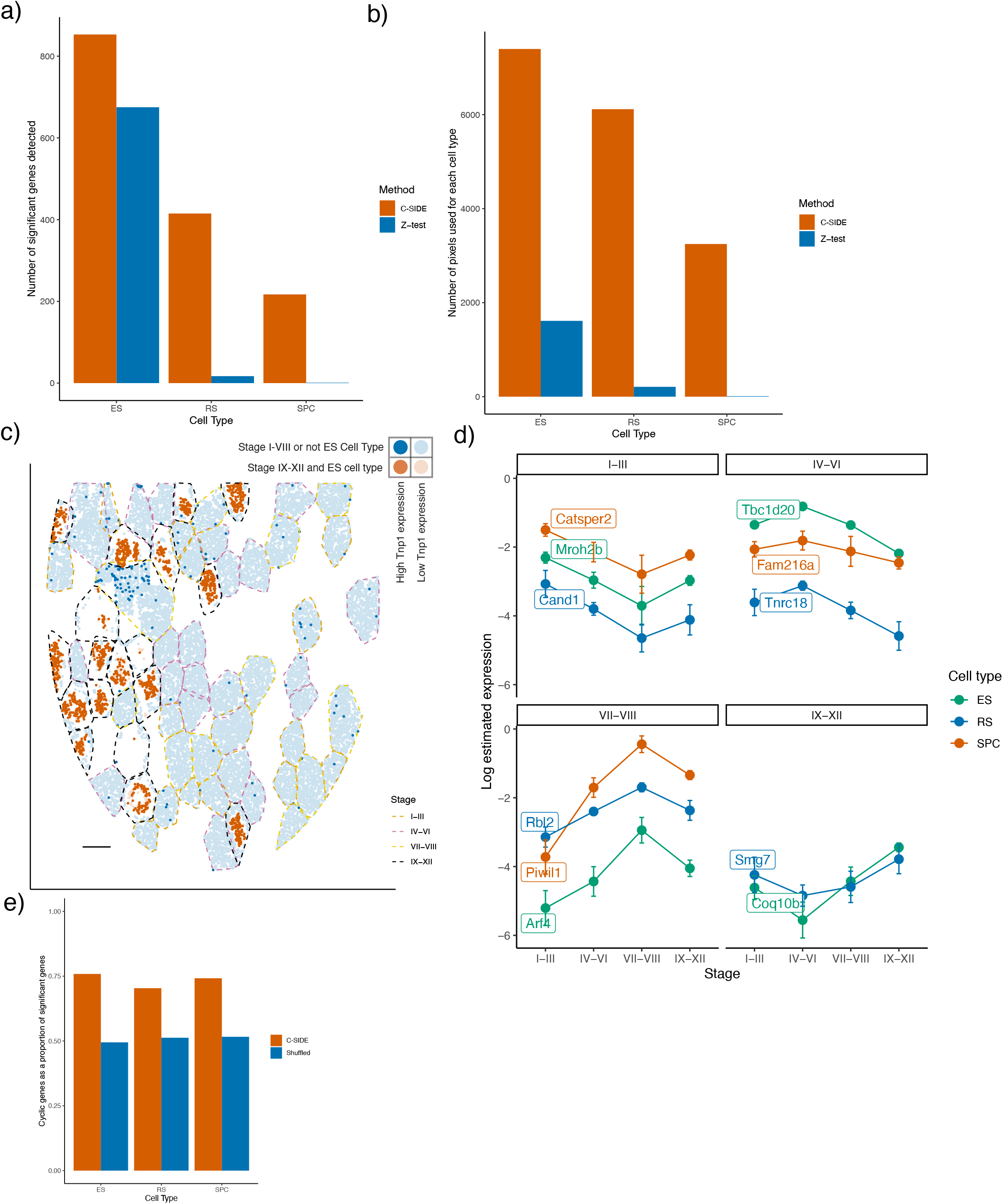
On the Slide-seq testes, C-SIDE achieves increased power in the presence of cell type mixtures to discover tubule stage-specific genes and cyclic genes. (a) Number of significant genes detected, for each cell type, by C-SIDE or the Z-test method. (b) Number of pixels used, for each cell type, to fit the C-SIDE or Z-test model. (c) Spatial plot of *Tnp1*, a gene identified by C-SIDE to be differentially expressed in stage IX-XII of cell type ES. Red represents the pixels of cell type ES within stage IX-XII, whereas blue represents pixels of another cell type or region. Bold points represent pixels expressing *Tnp1* at a level of at least 7.5 counts per 500. Scale bar represents 250 microns. (d) For each cell type, genes identified using C-SIDE results to be cyclic. Panels, indexed by tubule stage, contain cyclic genes whose peak estimated expression is at that stage. Error bars represent confidence intervals. (e) Proportion of genes categorized as cyclic (using C-SIDE fits), compared to proportion that would be categorized as cyclic if tubule stages were shuffled.

**Supplementary figure 5:**
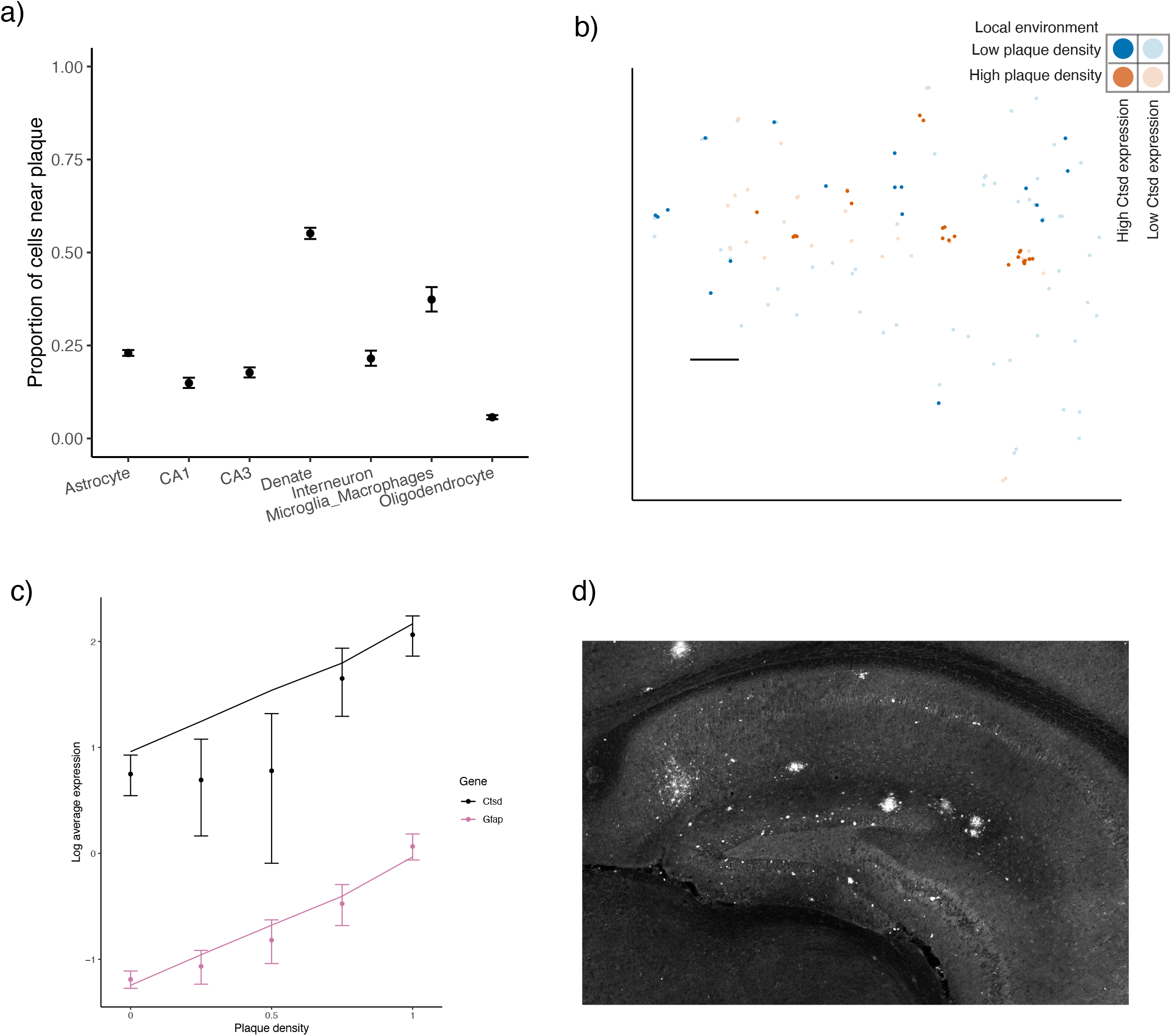
On the Slide-seq Alzheimer’s hippocampus, C-SIDE identifies genes whose expression depends on A*β* plaque density. (a) The proportion of cells, for each cell type, that localize in a high plaque density area. (b) Spatial visualization of *Ctsd*, whose expression within astrocytes was identified by C-SIDE to depend on plaque density. Red represents the astrocytes in high plaque density areas, whereas blue represents astrocytes in regions of low plaque density. Bold points represent astrocytes expressing *Ctsd* at a level of at least 3 counts per 500. Scale bar is 250 microns. (c) Log average expression of genes *Ctsd* and *Gfap*, which were identified to be significantly differ-entially expressed by C-SIDE for microglia/macrophages and astrocyte cell types, respectively. Single cell type pixels are binned according to plaque density, and points represent raw data averages while lines represents C-SIDE predictions and error bars around points represent ± 1.96 s.d. (*Supplementary Methods*). (d) Antibody stain of A*β* plaque in adjacent hippocampus section. This image is subsequently transformed to calculate a covariate for C-SIDE.

**Supplementary figure 6:**
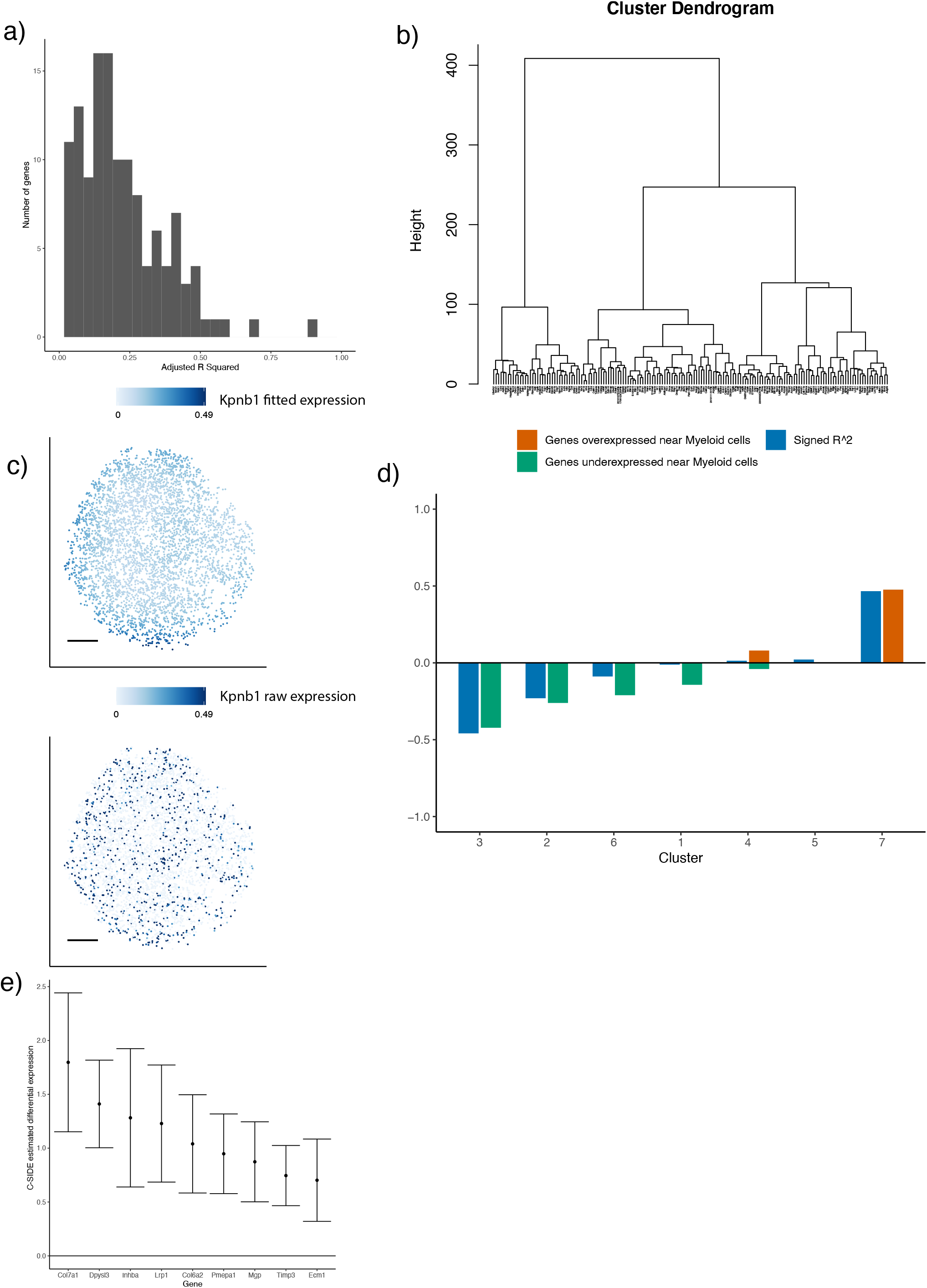
on the Slide-seq mouse tumor, C-SIDE identifies differentially expressed genes within tumor cells. (a) Histogram, across genes identified to be significantly DE within tumor cells by nonparametric C-SIDE, of adjusted *R*-squared, which is defined as the proportion of variance, not due to sampling noise, explained by the C-SIDE model. (b) Dendrogram of hierarchical clustering of (n = 162 significant genes) C-SIDE’s fitted smooth spatial patterns. (c) Spatial plot in tumor cells of *Kpnb1*, a *Myc*-target gene identified to be differentially expressed by nonparametric C-SIDE. Top shows C-SIDE fitted expression, while bottom shows observed expression in counts per 500. Scale bars are 250 microns. (d) For each cluster of spatially-varying genes, the proportion of genes identified by hypothesis-driven C-SIDE to be over- or under-expressed near myeloid cells. This proportion is plotted alongside the squared correlation of the cluster to the density of myeloid cells. (e) C-SIDE estimated differential expression and 95% confidence intervals of 9 genes from the epithelial-mesenchymal transition (EMT) pathway identified to be significant.

**Supplementary Figure 7:**
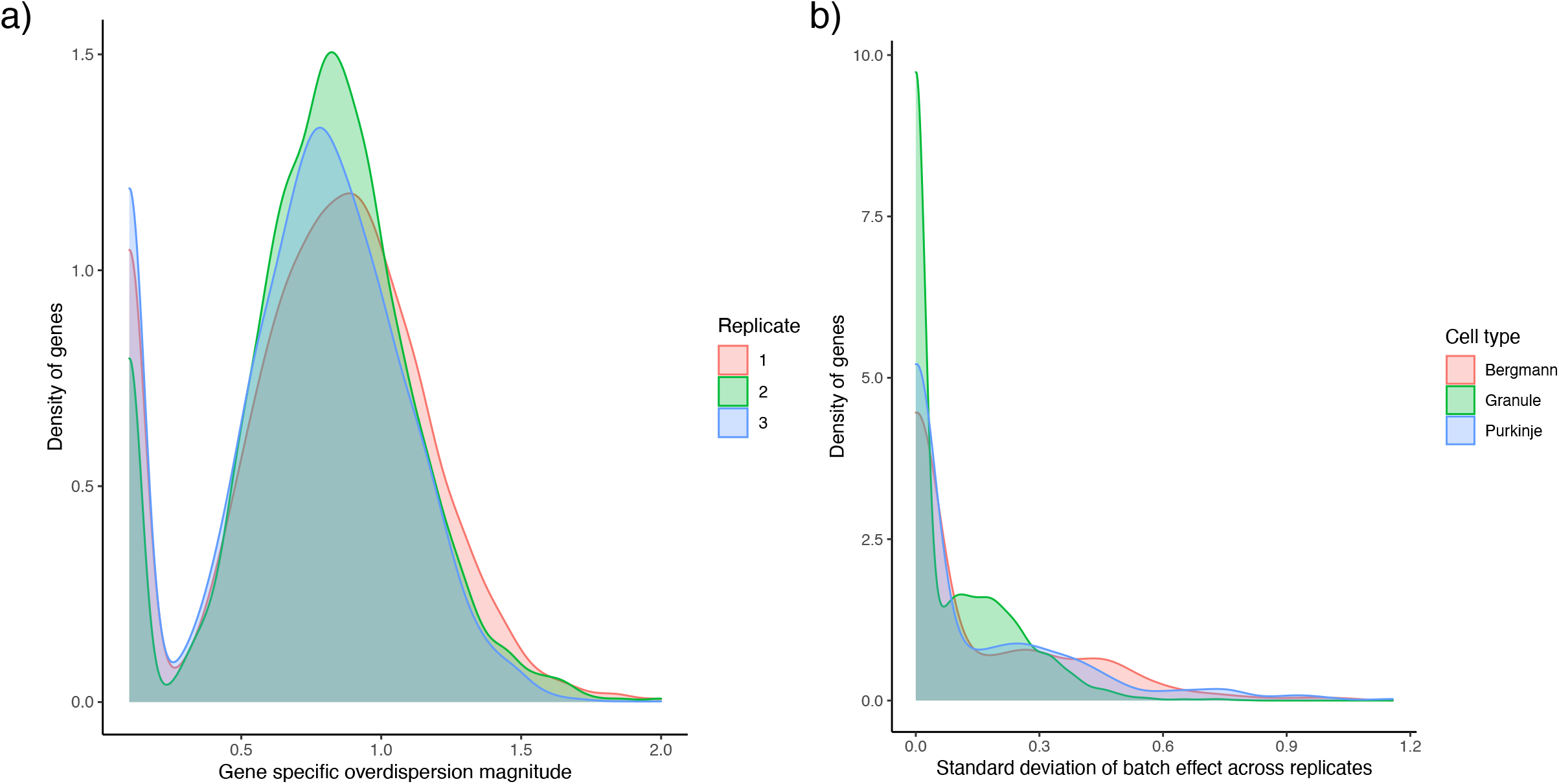
C-SIDE estimated variance parameters on the Slide-seq cerebellum data. (a) Density plot, over genes, of overdispersion standard deviation, *σ_ε_*, for each of three Slide-seq replicates. (b) Density plot, over genes, of C-SIDE estimated batch effect standard deviation, *τ*, for each of the Bergmann, granule, and Purkinje cerebellum cell types.

## Notes

https://www.github.com/dmcable/spacexr

### Competing Interest Statement

See conflict of interest statement in methods section

### Summary of Updates

Our method has been renamed as C-SIDE

